# Climate, caribou and human needs linked by analysis of Indigenous and scientific knowledge

**DOI:** 10.1101/2023.02.01.526335

**Authors:** Catherine A. Gagnon, Sandra Hamel, Don E. Russell, James Andre, Annie Buckle, David Haogak, Jessi Pascal, Esau Schafer, Todd Powell, Michael Y. Svoboda, Dominique Berteaux

## Abstract

Migratory tundra caribou are ecologically and culturally critical in the circumpolar North. However, they are declining almost everywhere in North America, likely due to natural variation exacerbated by climate change and human activities. Yet, the interconnectedness between climate, caribou, and human well-being has received little attention. To address this gap, we bridged Indigenous and scientific knowledge in a single model, using as example the Porcupine caribou herd social-ecological system. Our analysis, involving 688 (fall season) and 616 (spring season) interviews conducted over nine years with 405 (fall season) and 390 (spring season) Indigenous hunters from nine communities, demonstrates that environmental conditions, large-scale temporal changes associated with caribou demography, and cultural practices affect hunters’ capacity to meet their needs in caribou. Our quantitative approach bolsters our understanding of the complex relationships between ecosystems and human welfare in environments exposed to rapid climate change, and shows the benefits of long-term participatory research methods implemented by Indigenous and scientific partners.

## Introduction

Caribou and reindeer (both *Rangifer tarandus*, hereafter caribou) are among the most ecologically and culturally significant species roaming the circumpolar North. They shape ecosystems through their grazing, trampling, nutrient cycling, and support to multiple predators^1,2^. They also form tightly-coupled social-ecological systems (SES; an integrated perspective of humans-in-nature in which social and ecological subsystems are linked by mutual feedbacks, and are interdependent^3^) with many Indigenous communities across the Arctic and sub-Arctic, for whom caribou is both a key food source^4,5^ and a fundamental element of culture and identity^6,7^.

Since the beginning of this century, most North American caribou herds have declined dramatically^8,9^. Whereas Indigenous peoples and scientists alike have long documented the natural fluctuations of tundra populations (*R. t. groenlandicus* and *R. t. granti*)^10,11,12^, the current declines occur in a context of rapid climate change and growing human interference and industrialization, thus generating strong concerns for the ability of herds to rebound, for the sustainability of traditional Indigenous livelihoods, and for the overall health of northern ecosystems.

While the impacts of climate change on caribou populations are intensely studied^13,14^, the complex relations between climate, caribou, and the well-being of Indigenous peoples remain poorly understood. Difficulties in unravelling these relationships include lack of conceptual models capturing expected causal pathways, data gaps in critical variables, and the complexity of required statistical approaches^3,15^. Most importantly, fully assessing the context, motives and outcomes of Indigenous peoples’ relations to climate and caribou requires long-term partnerships with local communities and research endeavours that respectfully connect with Indigenous and local knowledge (ILK), here understood as “a cumulative body of knowledge, practice and belief, evolving and governed by adaptive processes and handed down and across (through) generations by cultural transmission, about the relationship of living beings (including humans) with one another and with their environment”^16,17^.

Disentangling the links between climate, the biosphere and human well-being is critical to advance sustainability research^18^. Demonstrating the biocultural consequences of climate and biosphere disruptions helps anchor global issues at the local scale^18^, and motivates monitoring and management practices that address the characteristics of subsistence resources while also accounting for the needs and perspectives of local users. Where concurrent changes in climate and land use are a looming menace for biodiversity and the traditional livelihoods of human communities^14,19^, interweaving ILK and scientific knowledge could improve our collective capacity to produce an enriched picture of complex systems^20-24^, while offering opportunities to engage in research that promotes dialogue, social justice, and Indigenous self-determination^21,25^. Yet, interweaving knowledge is fraught with challenges, inherent to the encounter of different cultures, worldviews, languages, priorities, and the power relations that have tainted these encounters for so long^20,21^. Considering the increasingly quantitative nature of environmental studies, there are also specific challenges to interweaving ILK and quantitative analysis in respectful ways^20,26^.

In this context, we sought to identify the mechanisms linking climate, caribou, and human capacity to satisfy cultural and subsistence needs in a human-caribou system offering a unique research environment. This system revolves around the Porcupine caribou herd (PCH; *R. t. granti*), ranging over 250,000 km^2^ in the Yukon (Canada), Northwest Territories (Canada), and northeastern Alaska (USA; Fig. 1 and Supplementary Methods). In this region, strong nutritional, cultural, and spiritual connections to caribou have been maintained over millennia by several Gwich’in, Inuvialuit, and Iñupiat communities and their ancestors^6,7,27^. Caribou, known as *vadzaih* (Gwich’in), *tuktu* (Inuvialuktun) and *tuttu* (Iñupiaq), remains a central component of local food systems^4,28^. The PCH is also currently one of the largest migratory tundra caribou populations in the World (ca. 218,500 individuals in 2017; ^29^), and the only North American herd that significantly increased in size since 2000^8^. Paradoxically, the PCH has also experienced one of the most acute climate changes on Earth, with annual temperatures raising by 3 to 3.5°C over its range between 1948 and 2016^30,31^. Finally, critical calving and post-calving PCH ranges lay within the coastal plain of the Arctic National Wildlife Refuge (Alaska), an area known to the Gwich’in people as “*Iizhik Gwats’an Gwandaii Goodlit*” (The Sacred Place Where Life Begins)^32^. Potential oil and gas development in the Arctic National Wildlife Refuge has been a source of fierce debates since the 1980s^33^. Plans by the U.S. administration to auction off leases for development in 2021 have generated a sense of urgency to further document the dynamics of the Porcupine caribou SES.

**Fig. 1.**
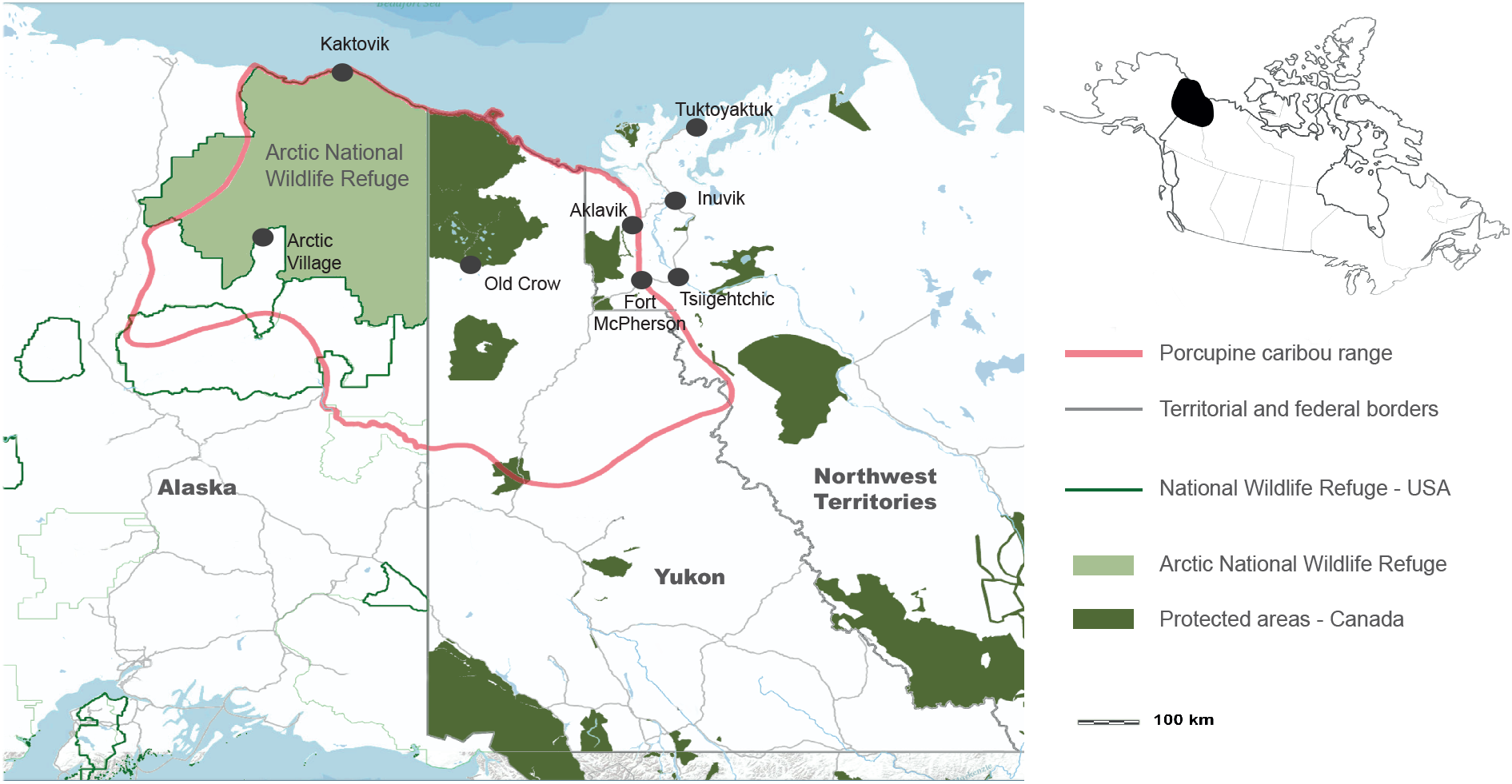
Annual range of the Porcupine Caribou Herd (PCH). Map showing the contour (pink) of the annual range of the PCH in northeastern Alaska (USA), and the Yukon and Northwest Territories (Canada). Indigenous communities that are connected to the PCH and that are part of the annual community-based monitoring program of the Arctic Borderlands Ecological Knowledge Society are identified with a black dot. Map data from <u>OpenStreetMap</u>.

Here, we developed a long-term collaboration between researchers and northern community members to design and implement a research approach to evaluate the general hypothesis (Fig. 2a) that climate influences the ability of Indigenous hunters to meet their needs in caribou. Previous studies drawing on scientific research and ILK have generated scenarios suggesting how climate warming could reduce caribou harvest in Porcupine caribou communities, through induced changes in land access^34,35^. We used a unique and comprehensive model bridging long-term scientific data and ILK to test the direct and indirect effects of regional temperature, snow conditions, icing events, and large-scale temporal changes associated with caribou demography on caribou distribution and the perceptions and behaviour of hunters (Fig. 2). We analysed 688 (fall season) and 616 (spring season) interviews conducted over nine years (2000-2008) with 405 (fall season) and 390 (spring season) Indigenous hunters from nine communities located within the PCH range (Fig. 1 and Methods). These interviews were performed through the community-based monitoring program of the Arctic Borderlands Ecological Knowledge Society (ABEKS), a participatory research program designed and implemented by Indigenous peoples and scientists since 1998^36,37^. We also used 7,428 caribou locations obtained via satellite monitoring of 32 adult females to estimate caribou distribution in relation to communities (hereafter referred to as caribou distribution). Using piecewise structural equation modelling (SEM)^38^, we then assessed the causal relationships between environmental conditions, time (years), caribou distribution in relation to communities, hunters’ perceptions of caribou availability, hunting activities and, ultimately, hunters’ capacity to meet their needs in caribou.

**Fig. 2.**
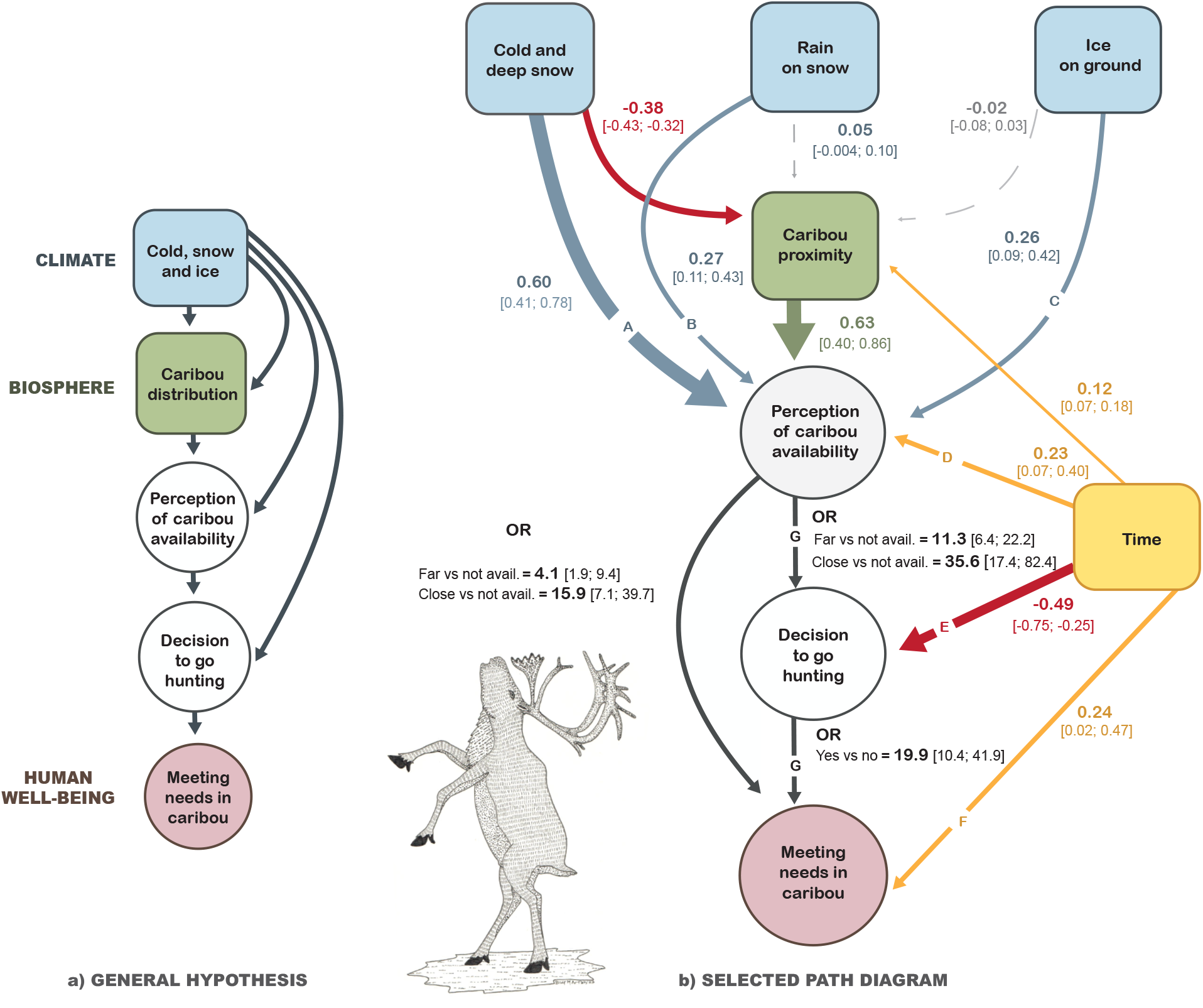
General hypothesis and selected path diagrams of the relationships between environmental conditions, time (years), caribou distribution in relation to communities (proximity), Indigenous hunters’ perceptions of caribou availability, hunting activities, and meeting needs. **a**, General hypothesis evaluated, which stipulates that climate influences the ability of Indigenous hunters to meet their needs in caribou (see Supplementary Tables 6 and 7 for the full list of hypothesized causal models). **b**, Final model testing how the capacity of hunters to meet their needs in caribou during the fall is directly and indirectly affected by environmental conditions and time. All continuous variables (in blue, green, and yellow) were standardized (Supplementary Table 8), meaning these parameter estimates (path coefficients) can be compared to assess their relative influence, with the width of the arrows scaled to the strength of the path coefficients. Solid arrows represent clear unidirectional relationships between variables (i.e. the 95% confidence interval (CI) excludes 0). Arrow colors, other than red, refer to the associated continuous variables and represent positive relationships. Red arrows emphasize negative relationships. Grey dotted arrows indicate lack of evidence for a clear relationship (95% CI overlapping 0). Path coefficients are presented as odds ratios (OR) when the response and the explanatory variable were both categorical. Blue and green squares indicate that data were generated through meteorological instruments and satellite collars. White and pink circles indicate that data were generated through interviews with hunters. Causal pathways lettered **A** to **G** correspond to the panels **a** to **g** in Fig. 3. Copyright for the caribou drawing: Inuit artist Billy Merkosak.

“Meeting needs” rests mainly on the idea of having enough caribou to meet subsistence requirements. However, it also relates to culture, identity, and spirituality, as well as to an individual assessment of fulfillment, influenced by a series of socioeconomic factors (e.g. family size). Therefore “meeting needs” cannot be assumed to correlate with a specific number of harvested caribou. For northern Indigenous peoples, being on the land and harvesting country food is intricately linked to well-being^7,39^. For Porcupine caribou hunters, caribou well-being also depends on how humans respect caribou^6,7^. The term Porcupine caribou SES refers to this nexus of complex cultural and ecological feedbacks between humans and caribou^40^. Here, we explore the climate-caribou-hunters loop to investigate the potential impacts of climate change on the subsistence livelihoods of Indigenous communities, a question of local to global relevance^20,41^. Insights learned from this study, both in terms of results and methods, should resonate for many other social-ecological systems.

## Results

### Effects of climate

As predicted by our conceptual model (Fig. 2a), we found that environmental conditions had direct influences on caribou distribution in relation to communities and hunters’ perceptions of caribou availability (Figs. 2B and 3, model fit: *C* = 27.98, *k* = 28, *P* = 0.47). Our model revealed that the composite variable “cold and deep snow”, describing snow depth, cold temperatures, and early snow arrival (Supplementary Tables 4 and 5), was the environmental variable with the strongest effect on both caribou distribution (Fig. 2b) and hunters’ perceptions of caribou availability (Fig. 2b, path A and Fig. 3a). Comparing the relative strength of path coefficients, this climate effect was ∼1.5 times stronger on hunter’s perception than on caribou distribution. Counterintuitively, colder falls with early and deep snow conditions corresponded to caribou being further away from communities in terms of distribution, but also to caribou being considered more available by hunters. Specifically, for an increase of 1 m of cumulated snow, caribou were ∼88 km further away from communities (Supplementary Fig. 2a), but hunters almost tripled their probability to consider caribou as being close to their community, and thus available (Fig. 3a).

**Fig. 3.**
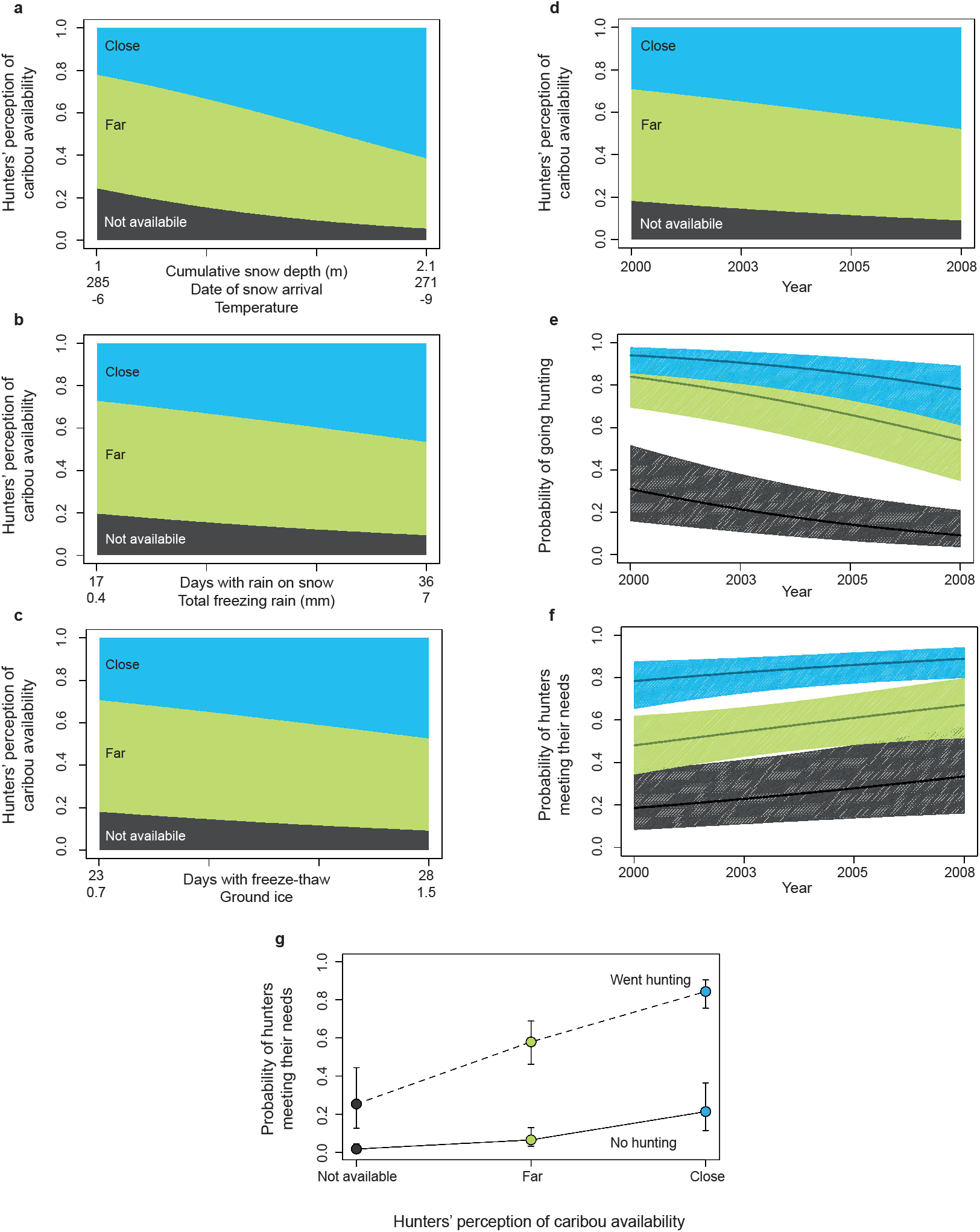
Effects of climate and time (years) on Indigenous hunters’ perceptions, hunting activities, and capacity to meet needs in caribou during fall. Panels **a** to **d** show the cumulative probabilities (proportion) for hunters to perceive caribou as being close (blue), far (green), or not available (grey), in relation to snow depth, date of snow arrival, and temperature (**a**), number of days with rain on snow and quantity of freezing rain (**b**), number of days with freeze-thaw events and quantity of ground ice (**c**), and years (**d**). For these four panels, an increase in the size of a coloured area represents an increase in proportion. In panels **a** to **c**, the figure presents the predictions based on the scores of the principal components (PC) used as indices of snow, temperature conditions, and icing events. Because PC scores are meaningless, we present the corresponding values for the variables represented by each PC (i.e. variables with eigenvectors higher than 0.5 for each PC axis; Supplementary Table 4). Panels **e** to **f** show the effects of time (years) on the probabilities for Indigenous hunters to go hunting (**e**), and to meeting their needs in caribou during the fall (**f**). Solid lines represent the estimated mean probability and coloured zones the 95% confidence intervals (CI), according to a specific class of perceived caribou availability. Panel **g** shows the relationships between hunters’ perception of caribou availability and the probability to meet their needs in caribou depending on whether or not they went hunting. Each dot represents the estimated mean probability and error bars represent the 95% Cis (n=688). Panels **a** to **g** also correspond to causal pathways lettered **A** to **G** in Fig. 2.

To a lesser extent, icing events also impacted caribou distribution and perception of caribou availability (Fig. 2b, paths B and C; Figs. 3b and c). The composite variable “rain on snow”, representing an increase in frequency of days with rain on snow and in total freezing rain (Supplementary Tables 4 and 5), had a slight positive effect on caribou distribution in relation to communities (Fig. 2b) and had a moderate positive effect on perception of caribou availability (Fig. 2b, path B and Fig. 3b). When frequency of rain on snow events increased from 17 to 36 days and total freezing rain increased from 0.4 to 7 mm, caribou were ∼12 km closer to communities (Supplementary Fig. 2b) and hunters almost doubled their probability to consider caribou as being available (Fig. 3b). Similarly, the composite variable “ice on ground”, representing an increase in days with freeze-thaw events and in ground ice thickness (Supplementary Tables 4 and 5), had a moderate positive effect on perception of caribou availability (Fig. 2b, path C and Fig. 3c). When frequency of freeze-thaw events increased from 23 to 28 days and ground ice thickness increased by 0.8 mm, hunters almost doubled their probability to consider caribou as being available (Fig. 3c).

Ultimately, climate impacted hunters’ decision to hunt or not and their capacity to meet their needs in caribou (Fig. 2b and 3) through indirect effects modulated by positive relationships between caribou distribution in relation to communities, perception of caribou availability, hunting activities, and meeting needs (Fig. 2b, paths G and Fig. 3g). For instance, hunters were ∼35 times more likely to go hunting when caribou were considered close versus not available (Fig. 2b, path G), which indirectly increased by ∼20 times their chance to meet their needs by going hunting (Fig. 2b, path G). When caribou were considered close, hunters also had ∼16 times more chances to directly meet their needs without going hunting (Fig. 2b), a disconnection between hunting and meeting needs that can be explained by the cultural tradition of sharing meat (see Discussion).

### Effects of time

From 2000 to 2008, time had a small but positive effect on caribou distribution in relation to communities (caribou were ∼31 km closer in 2008 compared to 2000; Fig. 2b and Supplementary Fig. 2c), as well as a moderate positive effect on perception of caribou availability (Fig. 2b, path D and Fig. 3d). Interestingly, although hunting became less frequent (Fig. 2b, path E and Fig. 3e), the capacity of hunters to meet their needs increased (see Discussion, Fig. 2b, path F and Fig. 3f). Moreover, aerial censuses of the PCH indicated that the population increased from 123,000 animals in 2001 to 169,000 in 2010, 197,000 in 2013, and 218,000 in 2017^29^. During our study period, time was thus highly correlated with an increase in caribou numbers.

## Discussion

Our study shows that meeting needs in caribou is directly and indirectly influenced by a complex interplay between environmental conditions, trends in caribou demography, and cultural traditions that would not have been unveiled without input from both Indigenous and scientific knowledge. Using structural equation modelling to bridge both knowledges, we show that cold temperatures and deep snow during fall have strong effects on both caribou distribution and hunters’ perception of caribou availability, which in turn influence the decision to go hunting and the capacity of meeting needs in caribou. Our analysis shows that meeting needs improved over time, likely due to a growing caribou population, and that the tradition of sharing caribou meat can compensate, in terms of meeting needs, for the decision not to go hunting. Taken together, our results identify specific mechanisms that could make the Porcupine caribou SES vulnerable to predicted increases in temperatures and delayed snow precipitations during fall. Our results also suggest that managing for a healthy caribou population and valuing the persistence of the cultural tradition of sharing could help alleviate some of the negative impacts of a warming climate.

Looking at the effects of environmental conditions on the Porcupine caribou SES, we found that temperature and snow conditions (depth and length of the snow season) were the environmental variables with the strongest causal effects on both caribou distribution and hunters’ perception of caribou availability. The concept of caribou availability, here, reflects ILK where resource availability is based on its abundance, distribution, and also the capacity for hunters to access harvest areas (accessibility)^42^. This explains that while coldest fall temperatures and deepest and earliest snows corresponded to caribou being further away from communities, caribou were nevertheless perceived by hunters as more available. Caribou distribution, here measured with collared female caribou, represents the location of caribou in relation to communities. Our results on caribou distribution are coherent with other analyses and Indigenous observations showing caribou distribution to be influenced by both snow depth and timing of fall snow arrival^6,43^. While caribou distribution certainly influences perception of caribou availability (Fig. 2b), as explained by Porcupine caribou hunters, the availability of caribou to communities is also related to caribou numbers and how easily they can be accessed by hunters^34^. In Porcupine caribou communities, hunters use snowmobiles and, according to them, poor snow cover during fall and early winter can prevent traveling or impede the capacity to locate caribou tracks, even when caribou are close to communities. Low snowfall during fall also closes some areas from hunting due to rough trail conditions^44^, which increases the risk of equipment failure and makes travel difficult for older hunters. Therefore, areas considered ‘close’ to the community in good snow conditions can be considered ‘far’ in poor snow conditions^44^.

Our results highlight that paying attention to resource access by harvesters avoids misleading conclusions when evaluating the impacts of environmental conditions on subsistence livelihoods. This is often hardly achievable without close partnerships with local knowledge holders and resources users. Had we considered caribou distribution as a proxy to caribou availability, we would have wrongly concluded that shallow snow depth, which made caribou closer to communities, had positive effects on the capacity of hunters to hunt and meet needs. Such detailed understanding of the functioning of SESs is critical to predict their fate from climate change scenarios^42^.

Icing events had positive effects on perception of caribou availability, but these effects were moderate when compared to those of cold temperatures and snow (Fig. 2b and Figs. 3b and c). This positive relationship could be related to how icing events influence the capacity of hunters to travel, but this deserves additional research, especially given that icing events are projected to increase over time^45^. Icing events also had a small but positive effect on caribou distribution in relation to communities. We hypothesize, based on discussions with local hunters, that caribou shift away from severe icing conditions that impede foraging, leading them to shelter in valleys, where they are closer to communities.

From 2000 to 2008, caribou became distributed slightly closer to communities and were perceived as increasingly available (Figs. 2B and 3D). The observed increase in the PCH population between 2001 and 2010 contributed to make caribou more available, likely explaining why the probability that hunters met their needs increased even though the probability of going hunting decreased during the same time period.

In many northern Indigenous societies, sharing of country food by successful hunters is an essential aspect of harvesting practices and ethics^6,46,47^. Although Gwich’in, Inuvialuit, and Iñupiat societies have strongly changed over the past century, country food sharing remains an important component of local food systems^7,47,48^, likely maximizing the overall well-being of the community while reinforcing social bounds^6,7^(see also Supplementary Methods). This cultural tradition is represented in our model by the direct effect of perception of caribou availability on meeting needs; even without hunting, people are indeed ∼16 times more likely to satisfy their needs when caribou are perceived as close versus not available (Fig. 2b). This explanation is based on qualitative answers provided by interviewees (see Methods) who regularly mentioned that while not going hunting, their needs were met through donations from other hunters.

Our unique model allowed us to quantify the effect of country food sharing on the capacity of meeting needs, showing that there is more sharing when caribou are considered highly available. Our results thus suggest that an accessible, healthy, and abundant caribou population could have positive impacts in the communities in terms of capacity to satisfy needs, through both caribou availability and caribou sharing. Note however that sharing is socially complex^7,46^ and socio-economic factors such as access to hunting equipment, maintenance of social ties, and persistence of key households, which account for a large percentage of country food procurements, are also important to maintain the sharing economy^47,48^.

The PCH is the only North American tundra caribou herd that increased in size during the last two decades^8^. This increase was linked to favorable spring conditions^49^ and limited regional resource development^19^. However, scientists and Indigenous hunters know that caribou populations go through 40-70-year cycles of abundance^10-12^. The PCH will thus likely enter a declining phase again, in a context where climate warming should induce degraded snow conditions^31^, and thus degraded traveling conditions for hunters. There is thus a serious risk that any factor accentuating the anticipated caribou decline might push the Porcupine caribou SES over a threshold where the capacity of hunters to meet their needs is compromised. Oil and gas development in the Arctic National Wildlife Refuge, for instance, was identified as a threat to the PCH^32,33^. Accordingly, future research should investigate the potential cumulative impacts of industrial development and climate warming, considering both their ecological and societal consequences. Food is very expensive in the North, and high poverty rates characterize many northern communities, thus access to a healthy caribou population strongly determines food security, health, and cultural dignity, all critical precursors of human well-being^5^.

The usefulness of any model rests on the choice of input variables. While our study highlights the effects of environmental conditions on hunting and need satisfaction, it does not address detailed socio-economic factors that may also influence hunting^35,44^ and food sharing^46-48^. This limits our ability to fully understand the complex mechanisms by which hunters integrate information about current conditions with their own experience, shared knowledge, and cultural practices. In addition, although the variable “years” is highly correlated to fluctuation in the size of the PCH, other potential temporal changes, for instance in socio-economic conditions, could also correlate to this variable and explain some underlying processes (see also the discussion about time-varying confounds in “Supplementary Results and discussion for the modelling of time”). This points to interesting future research questions investigating, through SEM, the relative influence of environmental conditions and socio-economic factors at play in this Porcupine caribou SES. Moreover, we analysed the fall and spring seasons in two distinct SEM models (see Supplementary Results and Discussion for the spring season) to avoid building an overly complicated model. This simplification assumed that hunting during one season did not impact hunting (or meeting needs) in the next season.

We favored a simple set of ILK indicators in our analysis, drawing on brief answers to three questions. Integrating culturally based knowledge that is qualitative, multi-causal, and holistic, within the reductionist nature of scientific knowledge can be hazardous^20,26^. Still, repeated observations based on intricate relationships with the environment are also one face of Indigenous knowledge^50^. In the ABEKS, the monitoring of Indigenous observations in such a succinct way resulted from a collective decision amongst both Indigenous representatives and scientists. The large spatiotemporal scale of analysed observations still provided unique insights while being firmly grounded in ILK. Meanwhile, the interpretation of results has been culturally contextualized through analysis of qualitative answers and results from previous studies^34,35,44^, and validated via meetings and discussions between researchers and local community members (see Methods). This case study demonstrates that when such an approach is elaborated as a process that allows enacting relational accountability^25^, i.e. working together on a long-term, respectful, and reciprocal dialogue between researchers and ILK holders, it can sincerely contribute to an improved collective understanding of SESs as well as of each other.

The Porcupine caribou SES is a typical case where Indigenous communities are tightly related to the land and other living beings (*dingii nin dzhii nan* in Gwich’in), from which they derive cultural, spiritual, and nutritional benefits, while also being exposed to intense climate warming. This SES also offers a unique context where local communities and scientists interacted and co-produced knowledge over many years, thereby allowing ILK and scientific knowledge to connect over a time scale rarely achieved elsewhere. This context has allowed us to demonstrate that at the local and regional scales, the capacity to satisfy human needs in relation to caribou depends on environmental conditions (temperature and snow), access to a healthy caribou population, and the persistence of cultural practices. Since regional management cannot tackle climate change or caribou population cycles, conservation of sensitive caribou habitat and the inter-generational transmission of the Indigenous cultural tradition of meat sharing should be priorities. So far, the northernmost regions of North America have been considered as a tragedy of open access in terms of mining and oil development^19^. Now, the balance between economic development and biodiversity conservation must be revisited to ensure that caribou SESs are maintained into sustainable pathways.

While our research approach is directly relevant to better understand other circumpolar caribou SESs, it also has broader implications. Indeed, the current global environmental crisis calls for collaborative efforts and the mobilization of all sources of valid knowledge to advance sustainable resource management at multiple scales^16,17^. In the context where global assessments look for innovative and transparent ways to bridge knowledge systems^16,41^, we have illustrated how bridging ILK and scientific data over the long term allows innovative analyses to identify mechanisms linking climate, biodiversity, culture, and human well-being. In culturally specific contexts, the involvement of relevant partners at all steps of the research process is particularly critical to understanding complex SESs^16,20,23^. Most importantly, long-term reciprocal partnerships can not only improve knowledge, but also advance communication and trust to create safe spaces where scientists, local resource users, and managers can negotiate research and management priorities^18,22,25^. Such inclusive processes may in turn allow a deeper access to the wisdom and experience of all partners and support the development of mutually acceptable management measures^23^ and, hopefully, more equitable futures^16^.

## Methods

This research complies with the ethical regulations of the Aurora Research Institute (Licences no. 13935, 14271 and 14989). Analyses of ABEKS’ interview data were also approved by each Indigenous community involved in the ABEKS’ project.

### Study area

Our study area encompassed the annual range of the PCH (Introduction and Fig. 1). Caribou have had considerable spiritual, cultural, and nutritional importance for Indigenous peoples of this region for thousands of years (Supplementary Methods)^6,40^. Within this range, the PCH undergoes bi-annual migrations from its spring and summer ranges located on the arctic coastal plain and adjacent mountains of Alaska and the northern Yukon Territory and Northwest Territories, to its winter range in the southern mountainous and forested habitats of northeastern Alaska, northern Yukon, and the Northwest Territories^51^. Since the 1970s, the PCH has received considerable attention due to potential industrial development over its calving range, as well as an observed population decline from 1989 to 2001^52^.

During this period the PCH declined from 178,000 to 123,000 individuals, but then increased to 169,000 individuals in 2010, 197,000 in 2013, and 218,500 individuals in 2017^52^. The PCH is harvested by the Indigenous communities of Old Crow, Fort McPherson, Tsiigehtchic, Aklavik, Inuvik, and Tuktoyaktuk, in Canada, and the Indigenous communities of Kaktovik and Arctic Village in the USA (Fig. 1). Culturally, these communities are Gwich’in (Old Crow, Fort McPherson, Tsiigehtchic, Aklavik, Inuvik, and Arctic Village), Inuvialuit (Aklavik, Inuvik, and Tuktoyaktuk) and Iñupiat (Kaktovik). As noted, two Indigenous groups, the Gwich’in and Inuvialuit, live in both Aklavik and Inuvik. Each of them was considered as a distinct community in the analyses.

### Data collected through interviews with hunters

Details about the creation, implementation, and guiding principles of the ABEKS have already been discussed in depth^36,37^. In short, ABEKS was created in 1994 out of a meeting between Indigenous leaders, community representatives, government managers, and researchers who wanted to improve ecological understanding within the range of the PCH by focusing on the respective strengths of both ILK and science-based research and monitoring. From the beginning, it was decided that ABEKS would be developed and managed cooperatively, and that control and ownership of the program and data would lay at the community and regional levels. As a non-profit society, ABEKS has an elected board of directors, but major decisions are to be taken by the broader membership. ABEKS directors and membership have varied over the years, but have always been composed by a majority of community representatives (e.g. community members; local Hunters and Trappers Committees) and regional organizations (e.g. Indigenous governments; wildlife co-management boards). One important aspect of the program is the ABEKS “annual gathering”, held each year in one of the participating communities or in the regional centers of Whitehorse and Inuvik. Annual gatherings allow participants to meet and exchange, to report on research and activities, to elect directors and to make important decisions such as the elaboration of an information-sharing protocol that respects community ownership of the data. During these gatherings, decisions are taken by consensus.

One of the core features of the ABEKS is also its annual community-based ecological monitoring program, which involves interviews with local experts from each PCH community, focusing on what the most active hunters, fishers, and berry pickers have observed over the preceding year. Since 1998, interviews are conducted each March by community monitors hired through local organizations (one monitor per community). A three-day training and planning session is held each year with the community monitors to practice interview techniques and review the program. One important task of community monitors is to identify, with the help of local organizations, a list of local experts to be interviewed. The target is 20 experts per community. Local experts are selected based on their knowledge, experience, and level of activity on the land during the previous year, including berry picking, hunting (various species) and fishing (Supplementary Methods). Prior to the interview, an overview of the program, and how the information will be stored and used, is presented by the community monitor. After this presentation, interviewees are invited to sign a consent form (see https://www.arcticborderlands.org/_files/ugd/ee3e9e_9698c702c3fd43188fc78c8c43d48593.pdf for the 2020 questionnaire and consent form). Interviews are confidential, but every participant is given a personal number that allows tracking answers from subsequent years without revealing identity.

Interviews are guided by a questionnaire (developed in 1997 and modified in 2003 and 2010) including both closed and open-ended questions on topics such as climate, caribou, berries, fish, and predators. Both the topics and format of the questions were designed out of a collaborative process between representative of Indigenous communities as well as governmental and university researchers. The reporting period for each interview includes previous winter, fall, and spring, meaning that an interview conducted in March 2008 covered the winter 2007-2008, fall 2007, and spring 2007 conditions. Once interviews are completed, community monitors are mandated to write a report that summarises interview results for their community. Each community monitor presents his/her summary at the annual gathering. Every summary report is then reviewed and validated by local organizations and compiled into a compendium coauthored by all community monitors. This compendium is distributed in communities, and a copy is sent to every interviewee. According to Eamer^36^: “This annual reporting by the community monitors to all contributors is crucial to the profile and success of the program. It allows people to see how their information is being used in developing a regional picture, and it reinforces community ownership of the results”.

This research is based on a long-term collaboration with ABEKS board of directors and community members that originated in 2008 during an annual ABEKS gathering. At that time, ABEKS had been conducting interviews for several years and had accumulated a large amount of data, but had done very little analyses with it. ABEKS’ directors wanted to move forward with analyses, but resources were limited. In 2008, there were also concerns that the PCH population was declining, and that climate change may have a role to play in this decline. C.A. Gagnon proposed to develop a collaboration to help with the analyses, trying to link climate data, caribou data, and the ability of hunters to meet their needs. This proposition was then identified as a priority. The ensuing collaboration was developed with the idea of ABEKS members and board of directors as partners in the research process, from the elaboration of ideas, to the interpretation and presentation of results. Over the years, this collaboration was maintained and reinforced via ongoing communication, attendance to the annual gatherings, community visits, attendance to the conference calls of the ABEKS board of directors, and individual conversations.

In accordance with the information-sharing protocol established by ABEKS, we had to obtain letters of support from every community participating in the ABEKS program as a first step to access ABEKS community-based monitoring data. In requesting letters of support, C.A. Gagnon explained how the data would be used, for what purpose, and how results would be reported back to communities. It was only after the letters were received that we could submit an official request for data access to the ABEKS board of directors, who then granted data access.

This study analysed answers to three questions relating to caribou availability, caribou hunting, and meeting needs in caribou (Supplementary Methods). In the first question analysed, interviewees were asked for each hunting season separately (i.e. fall, winter, and spring) how available caribou were to their community. Respondents had to choose between “close (within one day travelling distance, easily found)”, “far (within one week, required lots of efforts to get them)”, or “not available”. When caribou were hard to access for the community (i.e. “far”/”not available”), respondents were asked to explain what made them hardly accessible. Answers to these open-ended questions were provided in ca. 20% of the interviews. Thus, they could not be included in the quantitative analyses, but helped contextualize the analyses and interpret the results (see below).

In the second question, interviewees were asked whether they hunted or not in winter, fall, and spring. Only yes/no answers were allowed. If the answer was negative, hunters were asked to explain their answer. In the third question, interviewees were asked for each season whether they had enough caribou to meet their needs. Only yes/no answers were allowed. Interviewees were subsequently asked to explain their answer. Answers from this open-ended question, together with the published literature on food sharing as a mean to satisfy needs when not hunting, informed us on the importance of this cultural practice to explain results.

We analysed answers for the years 2000-2008, fall and spring periods. We chose to analyse data for spring and fall exclusively because they represent the most important hunting seasons. Obviously, preferred hunting seasons vary from one community to another, depending on their location in relation to the PCH migration routes. Nevertheless, the fall hunt is particularly appreciated because prior to the rut, caribou are considered “nice and fat”, and their hides are of high quality^6^. During fall and spring, caribou also gather in large congregations which offer interesting hunting opportunities^6^. Changes in the questionnaire prevented analysis beyond 2008. For the period covered by this analysis, between six and 22 local experts per community were interviewed each year, depending on the number of active knowledgeable experts identified by local organizations during a given year. There were no temporal trends in the number of hunters reporting in each community each year. For the fall, we analysed 688 interviews from 405 interviewees and nine communities. For the spring, we analysed 616 interviews from 390 interviewees and nine communities. Tuktoyaktuk was not included in the analysis due to the fact that they joined the program in 2004. Therefore, there was a gap in their data from 2000-2003.

Prior to any attempt at publication, we reported results to the annual gathering of the ABEKS which allowed communication with representatives from each community. Gathered participants reviewed results and assisted with interpretation and validation of findings. The participation of ABEKS board members (who are also community members) as co-authors to this article also helped ensure that the ideas and results presented in the paper are anchored in local realities.

### Climate data

Snow conditions have an impact on the movement and distribution of the PCH^43,53^, and on the distribution, winter survival, and feeding capacities of caribou in general^54^. Furthermore, icing events, i.e. climate events susceptible to create ice layers thick enough to impede access to forage (e.g. freezing rain), impact the distribution^55^ and population dynamics^45,55-58^ of caribou. To assess the influence of such climate conditions on caribou and their accessibility, we defined a series of seasonal variables based on climate data obtained through the CircumArctic Rangifer Monitoring and Assessment Network (CARMA^59^). The CARMA climate database^60^ was constructed using remotely sensed daily averaged climate data from the Modern Era Retrospective analysis for Research and Applications project (MERRA; https://gmao.gsfc.nasa.gov/reanalysis/MERRA/) and NASA (http://disc.sci.gsfc.nasa.gov/mdisc/data-holdings). These data have a spatial resolution of 1/2 degree (latitude) × 2/3 degree (longitude). To produce the CARMA database, shapefiles of known seasonal ranges for the PCH were overlapped with the MERRA gridded climate variables using ArcGIS version 10^61^. Daily data for a seasonal range was the median value among the grids included in the range polygons.

Daily averages specific to the fall (16 August–30 November), winter (1 December – 31 March), and spring (1 April–31 May) ranges of the PCH were then calculated^60,62^; Supplementary Table 1) to produce the seasonal climate variables (all variables are listed in Supplementary Tables 2 and 3). We calculated a series of variables describing snow conditions encountered by caribou over their seasonal ranges, as well as four variables describing the overall snow conditions for each year (number of days with snow, melt speed, melt date, and snow arrival date). We also calculated average temperatures (°C) for spring and fall. Finally, we calculated a series of variables describing icing events to which the PCH was likely exposed over its respective ranges. Similar variables describing icing events have been reported as importantly affecting caribou population fluctuations in the Arctic^57^. We used the term climate data to refer to any meteorological data used in our analysis.

Because climate variables were numerous and often correlated, we performed principal component analyses (PCAs; Supplementary Methods) to avoid multicollinearity and reduce the number of parameters used in models. Most importantly, it provided new sets of linearly independent variables, the principal components (PCs; Supplementary Tables 4 and 5) representing composite environmental variables (or “weather packages”^63^) describing year-to-year climatic variation that may better represent conditions experienced by living organisms.

### Caribou distribution data

The range use and migratory patterns of the PCH have been documented by U.S. and Canadian governmental agencies through satellite tracking of adult females since the 1980s (see details at http://www.pcmb.ca/herd). During 2000-2008, 32 cows were monitored for a total of 7,428 locations. The average duration of individual monitoring was 3.5 years, but some animals were followed for up to 9 years. Frequency of locations varied across seasons, years, and individuals. On average, individuals were located every 6 days (range 1-251), except from mid-May to mid-July, when locations occurred approximately every second day. Caribou location data were considered highly representative of the PCH location given that this herd remains strongly aggregated.

We analysed caribou and community location data using the sp^64,65^, rgdal^66^ and rgeos^67^ packages in R version 3.4.3. Locations were first imported in R and converted to the WGS84 coordinate system. We then calculated the weekly median distances between caribou locations and each of the 9 communities considered in our study. This allowed us to calculate the median of the weekly median distances for the fall (16 August-30 November) and spring (1 April-31 May) PCH seasons. Median seasonal distances between the PCH and the communities were highly correlated to minimum seasonal distances (r > 0.9). For ease of interpretation with indices of caribou availability, we report these values as the negative of median distances to represent the median distribution in relation to communities, representing caribou proximity.

### Piecewise Structural Equation Modelling

For each hunting season analysed (fall and spring), we performed a piecewise SEM^68^, also called confirmatory path analysis, to determine the relationships among: i-the different annual climate indices, ii-the study years, iii-the median distance between the PCH and communities, iv-the availability of caribou as perceived by hunters, v-the probability of hunters going hunting caribou, and vi-the probability that hunters met their needs. Piecewise SEM tests for direct and indirect causal relationships among variables in separate steps, one step for each endogenous variable^68^. Here, we analysed the climate-caribou-hunters’ causal relationships, ignoring the feedback loop of hunters’ effects on caribou. This approach was best suited to our study given the hierarchical and heterogeneous nature of our data^69^. Indeed, our dataset included continuous (climate, caribou distances, years), ordinal (caribou availability), and binary variables (probability of going hunting, probability of hunters meeting their needs), and piecewise SEM allows linking all these variables within one structural model^68^.

Piecewise SEM requires the *a priori* designation of plausible hypothesized relationships between independent and dependent variables, which are transposed in box-and-arrow diagrams called directed acyclic graphs^38,69,70^. To test the validity of these hypothesized causal models, simultaneous tests of all independence claims, known as a directional-separation (d-sep) test^69^, are performed. A directed acyclic graph model is rejected if the C statistics (i.e. Fisher’s C statistic, which will follow a chi-squared distribution if the data are generated under the causal hypothesis represented by the directed acyclic graph^66^) calculated from the d-sep test falls below the statistical significance level, set here at 0.05. We performed the piecewise SEM in five strategic steps (numbered 1 to 5 in Supplementary Fig. 3 and explained in details in Supplementary Methods), which illustrates our expectations of the most complex hypothesized causal structure linking fall variables. The modelling strategy consisted at testing alternative hypotheses based on different groups of variables at each step. Because variables included in the analysis were numerous, each step consisted of a SEM in itself and established the best reference model on which to add further variables to be tested in the next step. At each step of the piecewise SEM, we built a series of hypothesized models (Supplementary Tables 6 and 7) that we validated using the d-sep test^69^. We then selected the best model among the models considered valid based on the Akaike Information Criterion (AIC^71^). We considered the model with the lowest AIC as the best model unless other models were equivalent (Δ AIC ≤ 2), in which case we retained the most parsimonious model^72^. The selected model was then used as the basis model for the next step in which another level of variables was added.

For each season, the best model selected in the fifth step represented the complete final model of our piecewise SEM (fall: Fig. 2, spring: Supplementary Fig. 4 and Supplementary Results and Discussion). We calculated the path coefficients by regressing each variable on its direct causes ^69^ using the statistical models described in Supplementary Methods. For each path coefficient, we present the estimate and its 95% confidence intervals. All continuous variables were standardized to allow interpretation of the relative influence of different continuous predictors on a specific response^73^, meaning parameters presented are standardized (Supplementary Table 8). For models evaluating the causal links between perception of caribou availability (ordinal variable), decision to go hunting (binary variable), and meeting needs in caribou (binary variable), i.e. steps 4 and 5, coefficients were transformed as odds ratios when both the response and the predictor were categorical variables to ease interpretation and comparisons. Odds ratios measure effect size in logistic and multinomial regressions and range between 0 and infinity. Values around 1 indicate no difference, values moving from 1 towards 0 indicate a decreasing probability, and values moving from 1 to infinity indicate an increasing probability. As an example, an odds ratio of 2 indicates that an event is twice more likely to occur when the independent variable increases of 1 unit.

## Data availability

Climate variables used in this study and data on caribou distances to communities are available in the Dryad permanent repository^74^. Authors were not allowed to publicly archive survey data from the ABEKS due to their sensitive nature relating to endangered species and human identity. Access to ABEKS data requires consent from each Indigenous community involved in the project and the completion of a data request form which can be accessed via https://www.arcticborderlands.org/services.

## Code availability

All scripts used to run the analyses presented here are available in the Dryad permanent repository^74^.

## Acknowledgments

We thank the monitors, local experts, organizations, and community members who contributed to the Arctic Borderlands Ecological Knowledge Society since 1998. We thank N. Casajus for assistance in computing caribou location indices, and Y. Gendreau, V. Lamarre, P. Legagneux, P. Royer-Boutin, C. Doucet, L. Guéry, M.-H. Truchon, É. Bolduc, V. l’Hérault, and J. Bêty, N.G. Yoccoz and four reviewers for comments on the manuscript. This project was funded by the Canada Research Chairs Program 228343 (D.B.), the Natural Sciences and Engineering Research Council of Canada 85308851 (C.A.G. and D. B.), the Northern Scientific Training Program (Polar Knowledge Canada (C.A.G)), the International Polar Year program of Indian and Northern Affairs Canada 350391-07 (D.B), the Network of Centers of Excellence of Canada ArcticNet (D. B.), and the NSERC CREATE Training Program in Northern Environmental Sciences (Environorth, 370800-2010 (D.B.)). The community monitoring program of ABEKS was funded by the Government of Northwest Territories, the Khehłòk Eenjit Gwichít-Gwich’in Renewable Resources Board, the Government of Canada, the Indigenous Community Based Climate Monitoring Program, Parks Canada, Environment and Climate Change Canada, the North Yukon Renewable Resources Council, and the Wildlife Management Advisory Council (North Slope).

## Author Contributions

C.A.G., S.H., D.E.R, J.A., A.B., D.H., J.P. and E.S. conceived and designed the study. C.A.G and S.H performed the statistical analysis. C.A.G., S.H., D.E.R, J.A., A.B., D.H., J.P., E.S, M.Y.S., T.P and D.B. provided resources and analytical ideas. C.A.G, S.H, D.E.R and D.B. wrote the paper with substantial contributions from all authors.

