## Supplementary Information for "Climate, caribou and human needs linked by analysis of Indigenous and scientific knowledge"

### Supplementary Methods

#### The importance of caribou for Indigenous peoples

Caribou have had considerable spiritual, cultural, and nutritional importance for Indigenous peoples of this region for thousands of years[1,2]. Traditionally, caribou was an essential food source, but also provided material for clothing, tools, and artwork. In the Gwich'in culture, the creation story tells that long ago, caribou and humans were undifferentiated. As they separated, humans and caribou maintained a kinship relationship and agreed to become partners in the process of survival[2]. This partnership involved a set of norms for respecting caribou, including the rule to take only what one needs, and use all what is taken[3]. Today, caribou remains the most important source of adult daily protein, vitamin, and iron intake[4] in some of these communities, where food costs in market stores are higher than those found in southern Canada[5]. Harvesting opportunities, however, vary between communities according to their location in relation to the PCH annual migration. The PCH also continues to nourish a spiritual connection to the land: "The ability to procure one's own food and be able to share it with others is a source of satisfaction. Feelings of self-worth and dignity are closely related to personal skill and self-reliance. Sharing strengthens friendship and kinship bonds. A way of life centred around hunting as opposed to working for wages and buying imported store foods, is an integral part of northern society and culture"[6] (Supplementary Fig.1).

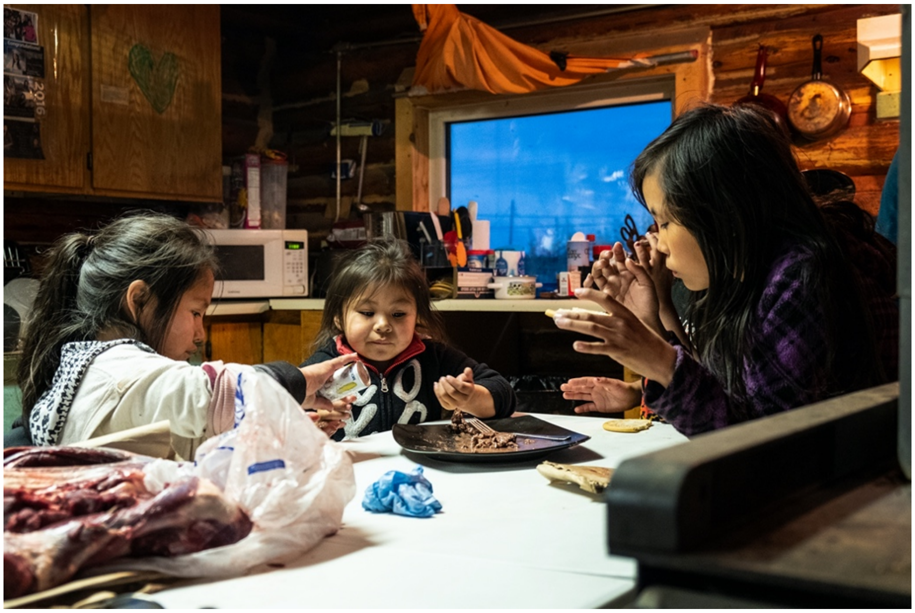

**Supplementary Fig. 1. Sharing a caribou meal with the younger generation.** Sharing caribou with children is not only a way to provide a nutrition meal, but also to strengthen social bonds and cultural transmission. Photo credit: Peter Mather.

**Questions analysed**

*Fall season*

1- How available were caribou to this community for hunting this past fall (until the beginning of the rut)?

   Choices of answers:
   a. close (within one day travelling distance, easily found)
   b. far (within one week, required lots of efforts to get them)
   c. not available

   Question follow-up:
   If it was difficult to get caribou this past fall, what made it hard?

2- Did you go caribou hunting last fall?

   Choices of answers:
   a. Yes
   b. No

   Question follow-up (if answer is no):
   What was the main reason that you didn't go caribou hunting last fall?

3- Did you get enough caribou last fall to meet your needs?
   Choices of answers:
   a. Yes
   b. No

   Additional notes on meeting or not meeting needs for caribou.

*Spring season*

1- How available were caribou to this community for hunting this past spring (April 1 – June 30))?

   Choices of answers:
   a. close (within one day travelling distance, easily found)
   b. far (within one week, required lots of efforts to get them)
   c. not available

   Question follow-up:
   If it was difficult to get caribou this past spring, what made it hard?

2- Did you go caribou hunting last spring?

   Choices of answers:
   a. Yes
   b. No

   Question follow-up (if answer is no):
   What was the main reason that you didn't go caribou hunting last spring?

3- Did you get enough caribou last spring to meet your needs?
   Choices of answers:
   a. Yes
   b. No

Additional notes on meeting or not meeting needs for caribou.

**Sample selection of key informants**

In most research involving ILK, the sampling strategy involves identifying key informants rather than selecting a random sample of informants in the community[7]. This is because interviews and questionnaires aim to collect information from people that are recognized by their community as having reliable and valid knowledge about the topic of interest[8].

In the case of ABEKS, the decision to interview “active knowledgeable experts” (i.e. purposive sampling) was made in the late 1990s, before the analysis presented in this paper was decided. The annual questionnaire aimed to gather information about a large array of topics related to the collection of living organisms, including berries, birds, fish, and caribou. The “active knowledge experts” were those community members most likely to provide reliable information about some of these topics. Due to the following three reasons, we considered the selection of “active knowledgeable experts” as a valid sample pertinent for our research questions, and we are confident that the external validity of the modeling results is not compromised by potential sampling biases.

First, experts were selected based on their activity and knowledge of the land in general (not just caribou hunting). This selection excluded community members with no experience on the land in a given year, thus ensuring the collection of valid information about environmental conditions. At the same time, the sampling included community members who were not necessarily the best caribou hunters, thus strongly decreasing the risk of excluding unsuccessful hunters whose needs might have been less likely to be met.

Second, the “active knowledgeable experts” involved in the ABEKS represented a large proportion (up to 17%) of the adult population of each community. Therefore, strong sampling biases were much less likely than if only a small proportion (e.g., <5%) was sampled.

Third, we included expert ID as a random effect in our models to reduce the effect of any remaining biases. This allowed the models to account for differences among hunters’ perceptions, reducing the chances of introducing bias in the results due to pseudo-replication if some hunters providing a unique set of information were more often interviewed than others.

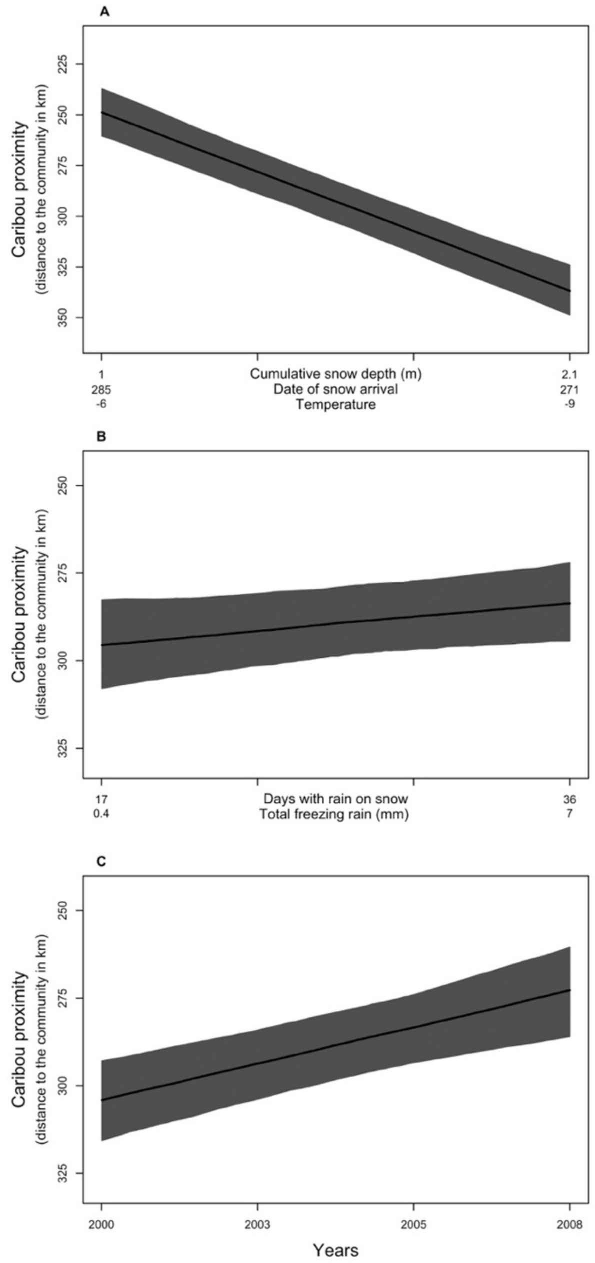

**Supplementary Fig. 2. Effects of climate and time (years) on caribou distribution in relation to communities during fall.** Caribou distribution in relation to snow depth, date of snow arrival, and temperature (A), number of days with rain on snow and quantity of freezing rain (B), and years (C). In panels A and B, the figure presents the predictions based on the scores of the principal components (PC) used as indices of snow, temperature conditions, and icing events. Because PC scores are meaningless, we present the corresponding values for the variables represented by each PC (i.e. variables with eigenvectors higher than 0.5 for each PC axis; Supplementary Table 4). Solid lines represent the estimated mean distance and grey zones the 95% confidence intervals (CI) predicted from the model (n=688).

**Principal component analyses**

Daily averages specific to the fall (16 August–30 November), winter (1 December – 31 March), and spring (1 April–31 May) ranges of the PCH were then calculated (Supplementary Table 1) to produce the seasonal climate variables (Supplementary Tables 2 and 3). Because climate variables were numerous and often correlated, we performed principal component analyses (PCAs) to avoid multicollinearity and reduce the number of parameters used in models.

PCAs generate new sets of linearly independent variables, the principal components (PCs[9]. The first PC accounts for the largest variation among input variables, and each successive PC accounts for the largest variation left in data once the previous PCs have been accounted for. Because we had numerous climate variables (Supplementary Tables 2 and 3) but few years recorded (n=9), we reduced the number of variables to ensure stability of the PCAs. First, we eliminated variables that were highly correlated (r > 0.7), and replaced them with one that provided similar climate information, such as average or cumulated snow depth (Supplementary Table 2). We then separated the remaining variables in two categories, those describing snow and temperature conditions and those describing icing events. We therefore performed two separate PCAs and repeated this procedure for each season, spring and fall (Supplementary Table 4). Although the normality assumption is not required for performing a PCA, one variable for the spring season (i.e. freezing rain falling on the ground in winter) was highly skewed and therefore transformed using the natural logarithm to improve the linearity of the relationships among the variables included in the PCA. For each PCA, we determined the number of PCs to be retained using the scree-test method, based on the rapid decrease in consecutive eigenvalues[10]. We then extracted the scores of the PCs selected and used them in further analyses as indices of snow and temperature conditions and indices of icing events.

For the fall season, we retained one PC to summarize snow and temperature conditions and two PCs to describe icing events (Supplementary Tables 4 and 5). For the spring season, we retained two PCs to summarize snow and temperature conditions and one to describe icing events (Supplementary Tables 4 and 5). For spring, since PCsnow1 was strongly correlated with years (time; r = 0.85), we only considered the variable PCsnow1 in the model, which therefore represented the combined effect of temporal changes and changes in snow and temperature conditions. Moreover, because we had to perform separate PCAs for snow and temperature conditions and icing events, PCs obtained from the separate PCAs were not necessarily independent. Nevertheless, their correlation was weak (r<0.4) and was accounted for in the piecewise structural equation modelling (SEM; see below). Although composite environmental variables generated by the PCA might make it more difficult to interpret causal links for each weather variable, caribou and hunters do not experience each environmental variable independently. On the contrary, composite variables (or “weather packages”[11]) may actually be experienced by living organisms and “often outperform local weather variables when it comes to explain climate-related variation in life history traits or animal numbers”[11].

**Supplementary Table 1.** Climate variables from the CircumArctic Rangifer Monitoring and Assessment Network (CARMA).

| CARMA climate variable | Unit | Calculation algorithm* |
|---|---|---|
| Mean daily snow depth | m | No algorithm. Equals to MERRA variable named as snow depth (snodp) |
| Mean daily temperature at 2 m above displacement | $^{o}$C | No algorithm. Equals to MERRA variable named as daily mean temperature at 2 m above displacement (t2m) |
| Mean daily fractional snow-covered area | fraction | No algorithm. Equals to MERRA variable named as fractional snow-covered area (frsno) |
| Number of days of freeze/thaw events | days | Cumulative days when t2m_max is above 0$^{o}$C and t2m_min is below 0$^{o}$C |
| Mean daily cumulative rain on snow | mm | Uses MERRA variables named as surface snowfall rate (precsno) and total surface precipitation rate (prectot). Is calculated as cumulative $rainfall_{(mm/s)}$*24*60*60 if (prectot-precsno)>0 & snodp >0.01 |
| Number of days of rain on snow | days | Cumulative days with rain on snow events. |
| Mean daily cumulative freezing rain | mm | Uses MERRA variables named as surface snowfall rate (precsno) and total surface precipitation rate (prectot). Is calculated as cumulative $rainfall_{(mm/s)}$*24*60*60 if (prectot-precsno)>0 & t2m <0 |
| Number of days of freezing rain | days | Cumulative days with freezing rain. |

List of climate variables calculated for each season from the CARMA database (SI Materials and Methods). *Calculation algorithms describe how CARMA variables were calculated from the Modern Era Retrospective analysis for Research and Applications (MERRA) daily averaged values.

**Supplementary Table 2.** Climate variables likely affecting the Porcupine Caribou Herd (PCH) over its fall range.

| Climate variable | Unit | Calculation | Season of analysis | PCH range* | Included in PCA |
|---|---|---|---|---|---|
| Average temperature–fall range | °C | Mean temperature for the fall season | Fall | Fall | Yes |
| Cumulative snow depth–fall range | m | Daily snow depth added over the fall season | Fall | Fall | Yes |
| Snow arrival date–fall range | Julian day | Julian day when snow cover reaches more than 20% on the fall range and never decrease below 20% again | Fall | Fall | Yes |
| Average snow depth–fall range | m | Mean daily snow depth for the fall season | Fall | Fall | No, correlated with cumulative snow depth – fall range† |
| Maximum snow depth–fall range | m | Highest daily snow depth recorded during the fall season | Fall | Fall | No, correlated with cumulative snow depth – fall range† |
| Variation in snow depth–fall range | | Coefficient of variation in snow depth for the fall season | Fall | Fall | No, correlated with cumulative snow depth – fall range† |
| Number of days with rain on snow–fall range | days | Number of days when rain on snow events occurred in the fall season | Fall | Fall | Yes |
| Number of days with freeze–thaw events–fall range | days | Number of days when freeze-thaw events occurred in the fall season | Fall | Fall | Yes |
| Number of days with freezing rain–fall range | days | Number of days when freezing rain events occurred in the fall season | Fall | Fall | Yes |
| Cumulative rain on snow–fall range | mm | Daily rain on snow added over the fall season | Fall | Fall | Yes |
| Cumulative freezing rain–fall range | mm | Daily freezing rain added over the fall season | Fall | Fall | Yes |
| Freezing rain falling on the ground–fall range | mm | Daily freezing rain when fraction of snow cover is less than 0.2, added over the fall season | Fall | Fall | Yes |

List of climate variables describing snow conditions, temperature, and icing events likely affecting the Porcupine Caribou Herd (PCH) over its fall range*, and calculated from CARMA data (Supplementary Table 1). *Variables were calculated with reference to the Porcupine Caribou Herd (PCH) seasonal range use. For the PCH, fall range is considered to be used from around 16 August to 30 November. † Variables highly correlated ($r > 0.7$) with another variable providing similar information were excluded from the principal component analysis (PCA) to ensure stability (see SI Materials and Methods).

**Supplementary Table 3.** Climate variables likely affecting the Porcupine Caribou Herd (PCH) over its spring range.

| Climate variable | Unit | Calculation | Season of analysis | PCH range* | Included in PCA† |
|---|---|---|---|---|---|
| Average temperature–spring range | °C | Mean temperature for the spring season | Spring | Spring | Yes |
| Cumulative snow depth–winter range | m | Daily snow depth added over the winter season | Spring | Winter‡ | Yes |
| Cumulative snow depth–spring range | m | Daily snow depth added over the spring season | Spring | Spring | Yes |
| Coefficient of variation in snow depth–spring range | | Coefficient of variation in snow depth for the spring season | Spring | Spring | Yes |
| Coefficient of variation in snow depth–winter range | | Coefficient of variation in snow depth for the winter season | Spring | Winter‡ | Yes |
| Melting date- spring range | Julian day | Julian day when snow cover reaches less than 20% on the spring range and never increase over 20% again | Spring | Spring | Yes |
| Average snow depth–spring range | m | Mean daily snow depth for the spring season | Spring | Spring | No, correlated with cum. snow depth–spring range† |
| Maximum snow depth–spring range | m | Highest daily snow depth recorded during the spring season | Spring | Spring | No, correlated with cum. snow depth–spring range† |
| Average snow depth–winter range | m | Mean daily snow depth for the winter season | Spring | Winter‡ | No, correlated with cum. snow depth–winter range† |
| Maximum snow depth–winter range | m | Highest daily snow depth recorded during winter | Spring | Winter‡ | No, correlated with cum. snow depth–winter range† |
| Number of days with freeze–thaw–spring range extended§ | days | Number of days when freeze-thaw events occurred on the spring range from 16 August to 31 May | Spring | Fall, winter and spring‡ | Yes |
| Number of days with freeze–thaw–winter range extended§ | days | Number of days when freeze-thaw events occurred on the winter range from 16 August to 31 Mars | Spring | Fall and winter‡ | Yes |
| Freezing rain falling on the ground–fall range | mm | Daily freezing rain when fraction of snow cover is less than 0.2, added over the fall season | Spring | Fall‡ | Yes |
| Freezing rain falling on the ground–spring range | mm | Daily freezing rain when fraction of snow cover is less than 0.2, added over the spring season | Spring | Spring | Yes |

| | | | | | |
|---|---|---|---|---|---|
| Freezing rain falling on the ground–winter range | mm | Daily freezing rain when fraction of snow cover is less than 0.2, added over the winter season | Spring | Winter‡ | Yes |
| Cumulative rain on snow–winter range extended§ | mm | Daily rain on snow falling on the winter range from 16 August to 31 March | Spring | Fall and winter‡ | Yes |
| Cumulative freezing rain–winter range extended§ | mm | Daily freezing rain falling on the winter range from 16 August to 31 Marc. | Spring | Fall and winter‡ | No, correlated with rain on snow–winter range expanded† |
| Cumulative rain on snow–spring range | mm | Daily rain on snow added over the spring season | Spring | Spring | No, correlated with rain on snow–winter range extended† |
| Cumulative freezing rain–spring range | mm | Daily freezing rain added over the spring season | Spring | Spring | No, correlated with rain on snow–winter range extended† |

List of climate variables describing snow conditions, temperature, and icing events likely affecting the Porcupine Caribou Herd (PCH) over its spring range*, and calculated from the CARMA database (Supplementary Table 1). *Variables were calculated with reference to the Porcupine Caribou Herd (PCH) seasonal range use. For the PCH, fall range is considered to be used from around 16 August to 30 November, the winter range is used from around 1 December to 31 March, and the spring range is used from around 1 April to 31 May. †Variables highly correlated ($r > 0.7$) with another variable providing similar information were excluded from the principal component analysis (PCA) to ensure stability (see Materials and Methods). ‡Climate variables describing conditions on the winter and fall ranges were included in spring analysis considering that caribou arriving on the spring range have been affected by climate conditions previously encountered. §Extended range means that the number of days icing or rain on snow events occurring on the spring or winter ranges were computed starting from the fall season. Again, caribou arriving on the spring range in April, for example, may encounter ice layers that were formed due to icing events occurring on the spring range during previous fall or winter.

**Supplementary Table 4.** Principal component (PC) scores (eigenvectors).

| | Snow and temperature | | Icing events | |
|---|---|---|---|---|
| Climate variable - fall | PC1 | | PC1 | PC2 |
| Average temperature – fall range (ºC) | **-0.60** | | | |
| Cumulative snow depth – fall range (m) | **0.56** | | | |
| Snow arrival date (Julian days) | **-0.56** | | | |
| Number of days with rain on snow – fall range | | | **0.50** | -0.38 |
| Number of days with freeze-thaw – fall range | | | 0.36 | 0.47 |
| Number of days with freezing rain – fall range | | | 0.39 | -0.22 |
| Cumulative rain on snow – fall range (mm) | | | 0.38 | **-0.52** |
| Cumulative freezing rain – fall range (mm) | | | 0.41 | 0.25 |
| Freezing rain falling on the ground – fall range (mm) | | | 0.40 | **0.50** |
| Climate variable - Spring | PC1 | PC2 | PC1 | |
| Average temperature – spring range (ºC) | 0.45 | 0.37 | | |
| Cumulative snow depth – winter range (m) | -0.37 | 0.36 | | |
| Cumulative snow depth – spring range (m) | -0.48 | 0.32 | | |
| Coefficient of variation in snow depth – spring range | 0.40 | -0.19 | | |
| Coefficient of variation in snow depth – winter range | -0.13 | **-0.75** | | |
| Melting date (Julian days) | **-0.51** | -0.20 | | |
| Number of days with freeze-thaw – spring range extended* | | | 0.49 | |
| Number of days with freeze-thaw – winter range extended* | | | **0.53** | |
| Freezing rain falling on the ground – fall range (mm) | | | 0.41 | |
| Freezing rain falling on the ground – spring range (mm) | | | 0.22 | |
| Freezing rain falling on the ground – winter range (mm) | | | **0.51** | |
| Cumulative rain on snow-winter range extended (mm)* | | | **0** | |

Principal component (PC) scores (eigenvectors) from principal component analyses (PCAs) including snow and temperature conditions, and icing events. These PCAs included climate variables likely affecting the Porcupine Caribou Herd over its fall and spring range (2000-2008). Numbers in bold identify variables with scores higher than 0.5 for each PC axis retained[63]. *Extended range means that the number of days of icing or rain on snow events occurring on the spring or winter ranges were computed starting from the fall season. Again, caribou arriving on the spring range in April, for example, may encounter ice layers that were formed due to icing events occurring on the spring range during previous fall or winter.

**Supplementary Table 5.** Principal components (PCs) selected for the path analyses.

| Component | Descriptive name of component* | Component's meaning | Season of analysis | % of variance explained | Cumulative % of variance explained |
|---|---|---|---|---|---|
| PC1– Snow and Temperature | Snow depth & Length of the cold snowing season | Greater PC scores represent colder falls with early snow arrival and more snow accumulation | Fall | 81.8 | 81.8 |
| PC1– Icing events | Frequency & intensity of all Icing events | Greater PC scores represent years with more icing events in general, particularly greater amounts of rain falling on snow | Fall | 42.2 | – |
| PC2– Icing events | Intensity of ground ice events (locked pastures) | Greater PC scores represent years with more freeze-thaw events and freezing rain falling directly on the ground (ground ice, forming ice locked pastures), but less frequent rain on snow | Fall | 29.8 | 72.0 |
| PC1– Snow and temperature† | Temperature & early melt | Greater PC scores represent years with a shorter snow season (early melting date), shallow snow in winter and spring, and warm temperatures | Spring | 56.5 | – |
| PC2– Snow and temperature | Snow depth | Greater PC scores represent years with less variability in the snow cover during winter, as well as deeper snow | Spring | 27.4 | 83.9 |
| PC1– Icing events | Frequency & intensity of all Icing events | Greater PC scores represent years with more icing events in general, particularly more freeze-thaw events and freezing-rain falling directly on the ground on the winter range | Spring | 47.7 | 47.7 |

Description of the principal components (PCs) selected to be used as climate variables in the path analyses, representing the snow and temperature conditions as well as the icing events to which the Porcupine Caribou Herd was exposed over its spring and fall ranges. *Names used in Fig. 2 or Supplementary Fig. 4. †This PC is highly correlated with years (2000-2008; *r* [95% confidence interval] = 0.85 [0.44; 0.97]).

**The five strategic steps of the piecewise SEM**

In the analyses, we included “years” as a continuous variable because we were interested in assessing whether there were temporal changes in this socio-ecological system. Temporal changes could have taken place because of the long-term temporal effects of changes in caribou demography or migratory patterns on their distribution, as well as the potential long-term changes in perception of fulfillment by hunters or in socio-economic factors influencing hunting capacities, which are not explained by annual variation in climate. We acknowledge that using the variable “years” could not identify the mechanisms behind potential temporal changes. In addition, the variable “years” could have been confounded by other unmeasured, latent variables, which are inherent to observational studies. Nonetheless, different piecewise SEMs provided similar results and conclusions (See Supplementary Results and Discussion) whether we modelled “years” as a fixed variable, fully excluded it, modelled it as a random intercept, or transformed the directed acyclic graph to a mixed acyclic graph (MAG) that allows to marginalize over potential latent variables[12]. The similarity of the results and conclusions suggests little influence of potential latent variables.

The first step of the piecewise SEM consisted of selecting the best model describing the causal structure among the different climate indices (PC scores relating to year-to-year climatic variation) as well as the continuous variable “years” (2000-2008). For the spring season, “years” (2000-2008) was highly correlated ($r$ [95% confidence interval] = 0.85 [0.44; 0.97]) with the PC describing temperature and melt date (PCsnow1). Therefore, “years” could not be included as a distinct variable in the modelling. Instead, temporal effects are confounded within the spring PC named “temperature and early melt”. PCsnow1 was therefore labelled “PCsnow1[years]” and should be considered as representing both large-scale temporal changes and temperature/melt date, with later years being warmer and having an earlier snowmelt. This first step of the piecewise SEM was based on the hypothesis that snow depth, the length of the snow season, and icing events could be linked via broader climate patterns, and that there may be trends in climate condition with time. Ten and six causal models were evaluated for the fall and spring (Supplementary Tables 6 and 7), respectively. Note that for each step, there are less models for the spring season because “years” and “PCsnow1” are confounded and are therefore never included in the same model as opposed to the fall season. This step was modelled with correlations or linear regressions using the lm function of R[13].

In the second step, we assessed the structural causal links between climate indices, time, and the median proximity between caribou and communities. This step is based on the hypothesis that snow depth, the length of the snow season, number of icing events, as well long-term processes (embedded with the variable “years”) influence caribou distribution. This hypothesis is based on previous work showing that environmental factors, especially snow conditions, influence PCH movements[14-16]. Studies performed on other migrating caribou herds also showed that icing events and demographic factors may influence caribou distribution[17]. Because environmental variables related to snow, temperature, and ice conditions have all been shown to influence caribou, we always considered all environmental variables together in the models evaluated. Therefore, the alternative hypotheses behind the models considered in this step had the following structure: i- a null model (the best model of the previous step), ii- a model including the effects of all environmental variables together, iii - a model including temporal changes only (years), and iv- a model including the effects of all environmental variables and temporal changes (years). This led to four causal models evaluated for the fall season (Supplementary Tables 6). For spring, however, we could only evaluate three models because the variable “PCsnow1[years]” represented temporal changes and environmental conditions, making it impossible to evaluate the second model representing the influence of environmental variables only (Supplementary Table 7). Note that the same logic applies to steps 3 to 5 to explain the lower number of causal models in the spring season. This step was assessed with linear mixed models (lmer function of the lme4 package in R[18]), including “community” as a random intercept to account for the repetition of caribou distances within a community each year.

In the third step, we evaluated the causal links between climate conditions, years, and caribou distribution in relation to communities, and hunters’ perception of caribou availability. Five and four causal models were evaluated for the fall and spring, respectively (Supplementary Tables 6 and 7). This hypothesis is based on informal discussions with hunters, combined with previous studies[19] showing that hunters have a deep and complex knowledge about how environmental conditions influence both caribou distribution and availability to hunters. To hunters, caribou availability embeds both the environmental conditions affecting caribou distribution and the capacity of hunters to access caribou[19]. We thus postulated that climate conditions and caribou distance to communities have both a direct effect on the hunter's perception of caribou availability (Supplementary Fig. 3). Because hunter perception of caribou availability was an ordinal variable with 3 levels ( "close", "far", "not available"), this step was assessed with cumulative link mixed models (clmm function of the ordinal package in R[20]) using a logit link and including "interviewee identity" nested within "community" as a random intercept.

In the fourth step, we assessed the structural links between climate conditions, years, perception of caribou availability, and whether or not local experts went hunting. Five and four structural models were evaluated for the fall and spring, respectively (Supplementary Tables 6 and 7). Again, informal discussions and previous research have shown that climate affects the probability that northern hunters go hunting and access wildlife[19,21]. In addition to climate, other environmental and socio-economic conditions (e.g. caribou density and access to gear) may also affect the probability of going hunting[22]. This step was modelled with generalized linear mixed models (GLMMs; glmer function of the lme4 package in R[18]) using a binomial family with a logit link and including "interviewee identity" nested within "community" as a random intercept.

In the fifth step, we tested the hypothesis that the probability that local experts meet their needs in caribou is directly affected by the probability that local experts go hunting, and directly or indirectly affected by climate indices, temporal trends in environmental and socio-economic factors, as well as by the perceived availability of caribou. Nine and six structural models were evaluated for the fall and spring, respectively (Supplementary Tables 6 and 7). This hypothesis is based on the fact that community members meet their needs in caribou both by going hunting themselves and through sharing of harvest within the community (see also [22]). This step was modelled with GLMMs as described in step four.

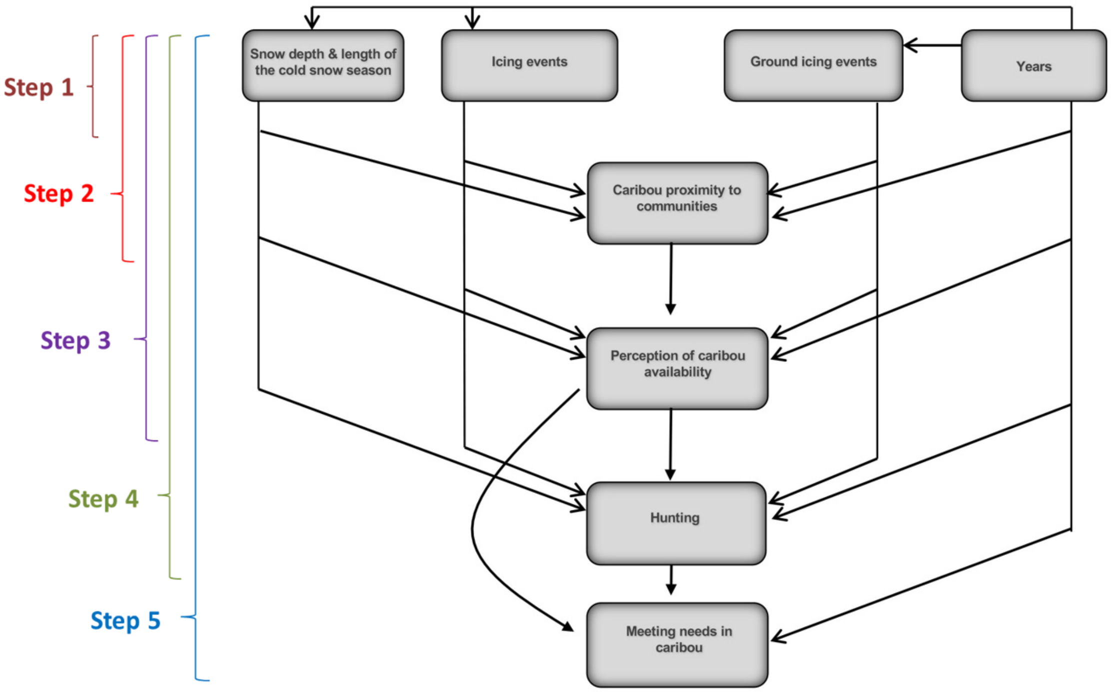

**Supplementary Fig. 3. Hypothesized path diagram.** Path diagram showing the most complex hypothesized causal structure for the fall season, linking direct and indirect relationships among climate conditions (indices obtained from PCAs), large-scale temporal variation (years), caribou distribution in relation to communities (reverse of median distance to communities), caribou availability as perceived by hunters, the probability of going hunting, and the probability that hunters will meet their needs, for the Porcupine Caribou Herd, northeastern Alaska (USA) and Northern Yukon and Northwest Territories (Canada), 2000-2008. The 5 steps illustrate the variables that were included in the 5 consecutive confirmatory path analyses that were conducted to build and select the final model.

### Supplementary Results and Discussion

#### Results and discussion for the spring season

For the spring season, we found that climate conditions and temporal trends had effects on caribou distribution in relation to communities, perception of caribou availability, hunting, and meeting needs in caribou (Supplementary Fig. 4). Similar to what was observed during fall, snow depth was the climate variable that had the greatest effect on caribou distribution and perception of caribou availability. When snow depth was deeper in spring, caribou tended to be further away from communities, but hunters tended to consider caribou as being more available. As was the case for the fall season, this can be explained by the fact that perception of caribou availability results from both caribou distance and hunters' traveling capacity. Hunters need snow to travel with snowmobiles, hence poor snow cover prevents access to caribou even when they are relatively close to communities. Hunters' ability to locate and access caribou may also be favoured by deeper snow conditions. Indeed, deep snow conditions may restrict caribou to shallow snow areas and slow down caribou movements[1,23,24]. Deep snow may also enhance tracking capacities by making footprints easier to detect. During spring, snow conditions had strong direct effects on the decision to go hunting or not. Whereas deep snow positively influenced the decision to go hunting, high spring temperatures and early melts had strong negative effects on hunting.

During the spring, there was a very strong correlation ($r = 0.85$) between years (2000 to 2008) and the principal component index related to temperature and early melts, meaning that average spring temperatures have increased from 2000 to 2008 and melting dates have tended to be earlier[25]. During the same time period, the Porcupine caribou population increased from 123,000 animals in 2001 to 169,000 in 2010. Consequently, for the spring season, it was not possible to separate the effects of climate and large-scale temporal effects that could be due to changes in caribou demography on caribou distribution, perception of caribou availability, hunting, and meeting needs. Nonetheless, the results show that temperature, early melt, and time (years) had a positive effect on caribou distribution, a weak positive effect on perception of caribou availability, a negative effect on hunting, and yet a positive effect on meeting needs (Supplementary Fig. 4). These effects are similar to what was observed for the large-scale temporal effects during the fall (Figs. 2 and 3). This leads us to believe that non-climatic temporal trends (such as caribou demographic trends) are more likely to explain the positive impact on meeting needs; if caribou were more abundant over time, they were more likely to be accessible to communities, and hunters could reduce hunting activities and yet increase their capacity to meet their needs (see main text).

Results also show that, in spring, icing events had a marginal positive effect on caribou distribution, but a strong negative effect on hunting (Supplementary Fig. 4). The negative effect of icing events on hunting was not observed in fall (Fig. 2). During fall, caribou is the main resource hunted by communities, whereas other resources, such as fish, are available in spring. Therefore, hunters may be less willing to go hunting during spring when climate conditions are not optimal. Nevertheless, if hunters decide to go hunting in years with more icing events, they are more likely to meet their needs in caribou than in years with less icing. This positive influence of icing events on meeting needs may be explained by the fact that icing events are producing a crust on top of snow which, according to Gwich'in Elders, makes travel on deep snow harder for caribou[1], likely resulting in higher success for hunters who decided to go hunting.

Similarly, as in the fall season, caribou distribution in relation to communities had a positive effect on perception of caribou availability (Supplementary Fig. 4). Hunters were ~28 times more likely to go hunting when caribou were perceived as being close versus not available, and they were ~29 times more likely to meet their needs when they went hunting. Furthermore, hunters were ~33 times more likely to meet their needs in caribou when they considered caribou as being close versus not available, even though they did not go hunting (Supplementary Fig. 4). As for the fall season, this paradox can be explained by the sharing tradition of caribou meat within the community, which seems to be enhanced when caribou are more available (Main text).

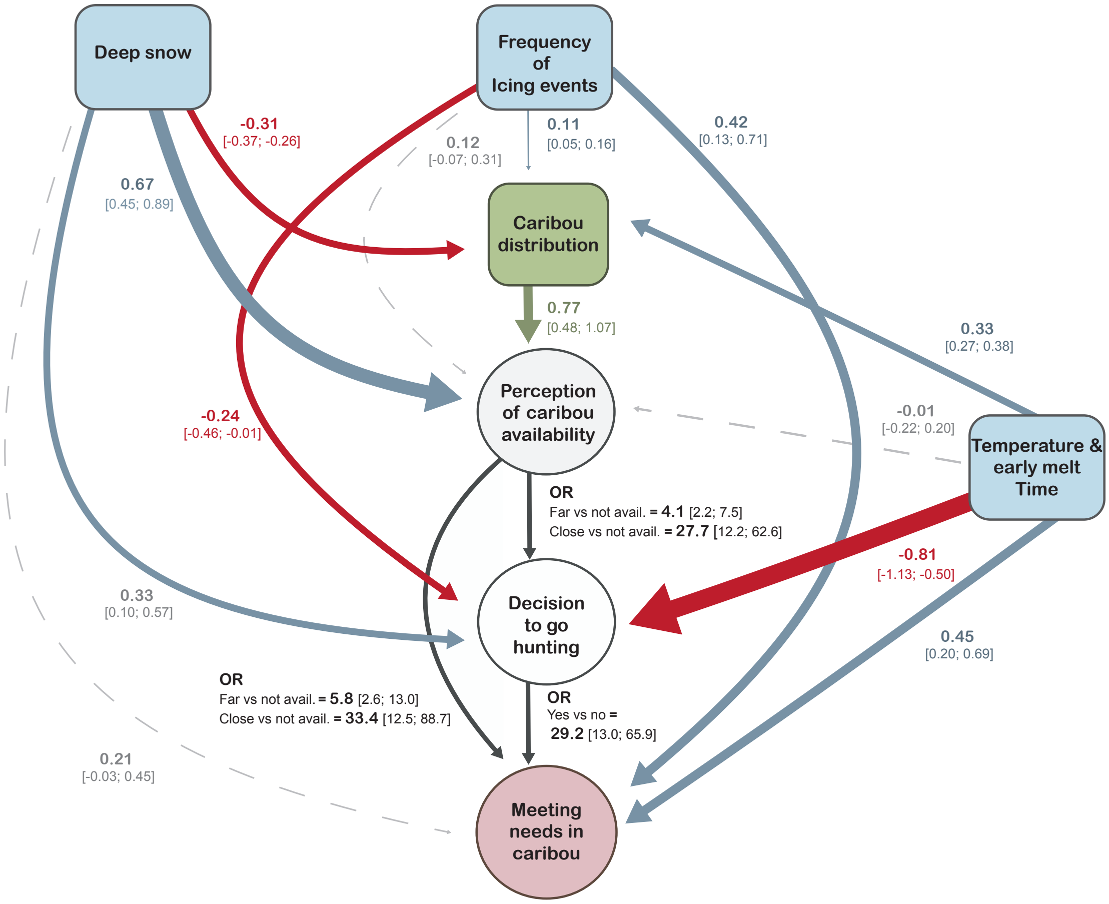

**Supplementary Fig. 4. Final model testing how the capacity of hunters to meet their needs in caribou during spring is directly and indirectly affected by environmental conditions and time.** Path diagram of the relationships between environmental conditions, time (years), caribou distribution in relation to communities (proximity to communities), Indigenous hunters' perceptions of caribou availability, hunting activities, and meeting needs (model fit: C = 6.78, k = 30, P = 0.8). All continuous variables (in blue and green) were standardized (Supplementary Table 8), meaning these parameters estimates (path coefficients) can be compared to assess their relative influence, with the width of the arrows scaled to the strength of the path coefficients. Solid arrows represent clear unidirectional relationships between variables (i.e. the 95% confidence interval (CI) excludes 0). Arrow colors, other than red, refer to the associated continuous variables and represent positive relationships. Red arrows emphasize negative relationships. Grey dotted arrows indicate lack of evidence for a clear relationship (95% CI overlapping 0). Path coefficients are presented as odds ratios (OR) when the response and the explanatory variable where both categorical. Blue and green squares indicate that data were generated through meteorological instruments and satellite collars. White and pink circles indicate that data were generated through interviews with hunters.

**Results and discussion for the modelling of time**

For the fall season (main text), we modelled the variable "years" as a continuous fixed effect because we were interested in estimating the temporal trends. For the spring season (Supplementary, p.15), we included the variable "temperature and early melt" as a fixed effect because this variable was strongly correlated with "years" and therefore represented the combined effect of temporal changes and changes in snow and temperature conditions. In both cases, there was a possibility for time-varying confounds due to unmeasured factors that vary among years. These time-varying confounds could bias the estimated parameters and thus the causal identification. Below, we present three additional models and contrast their results with those of our first model described above to evaluate the possibility of time-varying confounds in our system.

The second model fully excluded the variable "years", which meant for the spring season that we excluded both the variable "years" and "temperature and early melt". The third model included "years" as a random intercept. The fourth model was a transformation of the first model to a mixed acyclic graph (MAG). A MAG is an extension of a directed acyclic graph that allows to marginalize over potential latent variables (see Shipley & Douma 2021[12]). For that model, we used the function "toMAG" of the R package dagitty[26] to transform the first model to a MAG by including a single latent variable on all parameters. Therefore, that fourth model allowed us to estimate the parameters of the system with the underlying assumption that a time-varying confound was present on all variables in the system.

The results of these three alternative modelling approaches are compared to those of our first model in Supplementary Fig. 5 and 6 for the fall and spring seasons, respectively. We can clearly see that almost all parameters were similar in their means and 95% confidence intervals (CI). The parameters of the first model, including "years" as a fixed effect, and those of the second model, excluding this effect, were similar, demonstrating that including years as a temporal trend did not affect the other parameters estimated. The third model, which included "years" as a random intercept, provided similar mean estimates but wider CI for the weather variables. This was expected because of the study design. Indeed, all weather variables were averaged annual values, meaning there was only one value for each weather variable each year. Having "years" as a random intercept resulted in a model for which the fixed and the random effects were competing for the same information, which can lead to error-variance inflation[27] Although the third model provided less precise estimates, the overall results from that model offered conclusions similar to those of the first model.

Finally, the fourth (MAG) model also provided similar parameter estimates and CI to the first model. For that model, new paths were added in the transformation to a MAG. We can see these additional paths in the figures by looking at parameters for which there is only a purple estimate presented for a variable, e.g. the boxes representing parameter estimates for the 2$^{nd}$, 3$^{rd}$, 4$^{th}$ and 6$^{th}$ variables in Supplementary Fig. 5c. These represented additional paths that were possible under the assumption that a time-varying confound was present on all variables in the system. Almost all of these additional paths were relatively trivial (the CI of these estimates were overlapping 0), except for a few cases, i.e. some paths going from caribou proximity to hunting and meeting needs. These paths in the MAG mean that caribou proximity is a causal ancestor of hunting and meeting needs under the assumptions of this model, and that the statistical association between them, i.e. the apparent causal effect, could not be removed by statistically conditioning on any combination of the observed variables in the MAG (see Shipley 2016 and Shipley & Douma 2021 for more details on these paths that they referred to as "inducing paths"[12,28]). Even with these additional paths, this model provided similar results and similar conclusions, again suggesting that there was little influence of potential time-varying confounds in the system.

**a**. Effects on caribou proximity (step II, fall season)

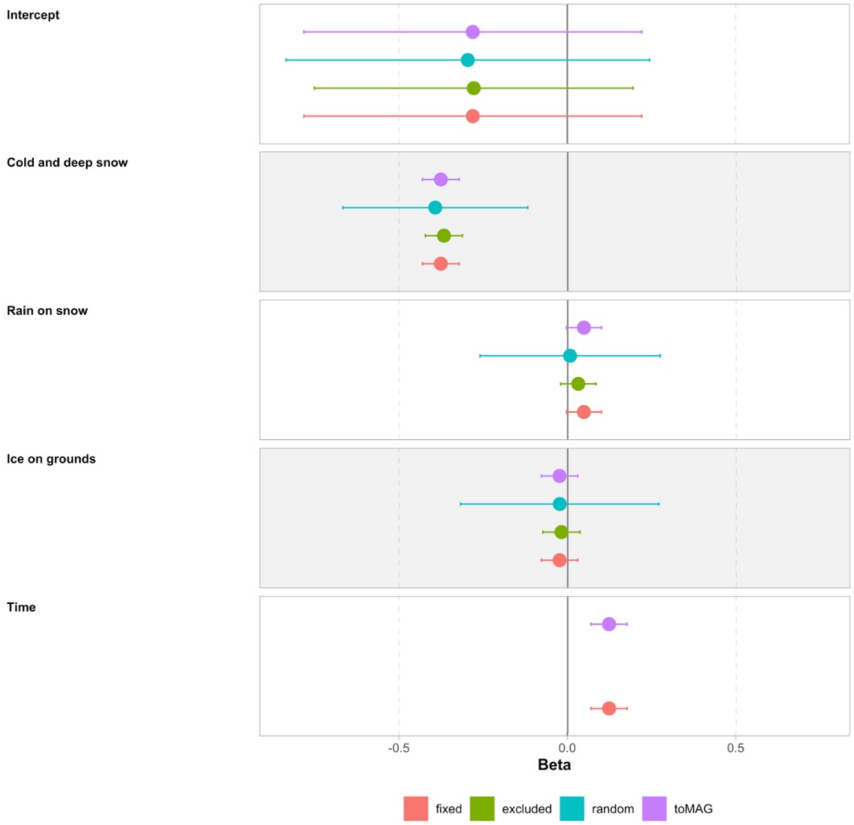

**Supplementary Fig. 5.** Mean parameter estimates (dots) and 95% confidence intervals (whiskers) contrasting four different methods to include the time effect “years” for the piecewise SEM of the fall season (n=688). The model presented in the main text is in red. This is a model including “years” as a fixed effect. In green is the model fully excluding “years”. In blue is the model including the variable “years” as a random intercept. In purple is the model transforming the model including “years” as a fixed effect to a MAG.

**b**. Effects on perception of caribou availability (step III, fall season)

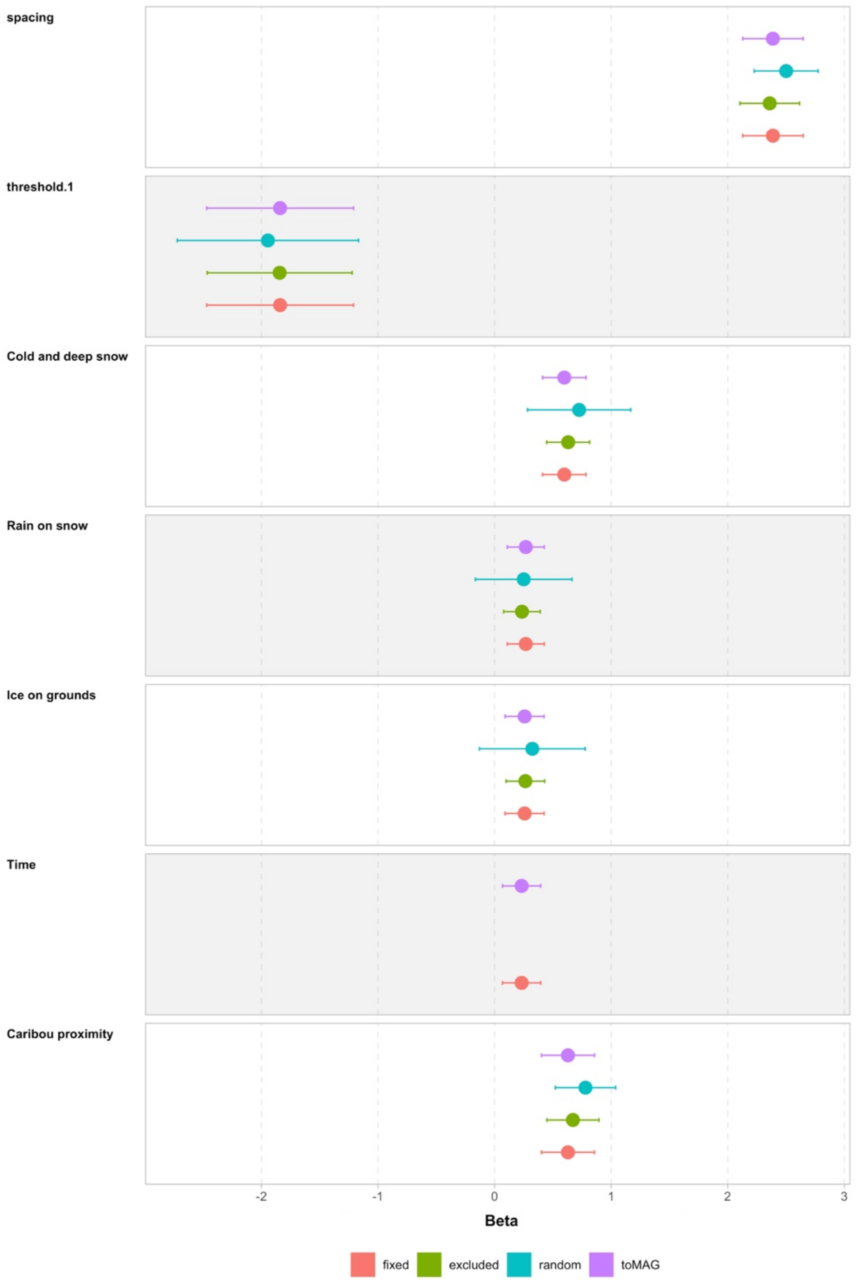

**Supplementary Fig. 5.** continued.

**c**. Effects on probability of going hunting (step IV, fall season)

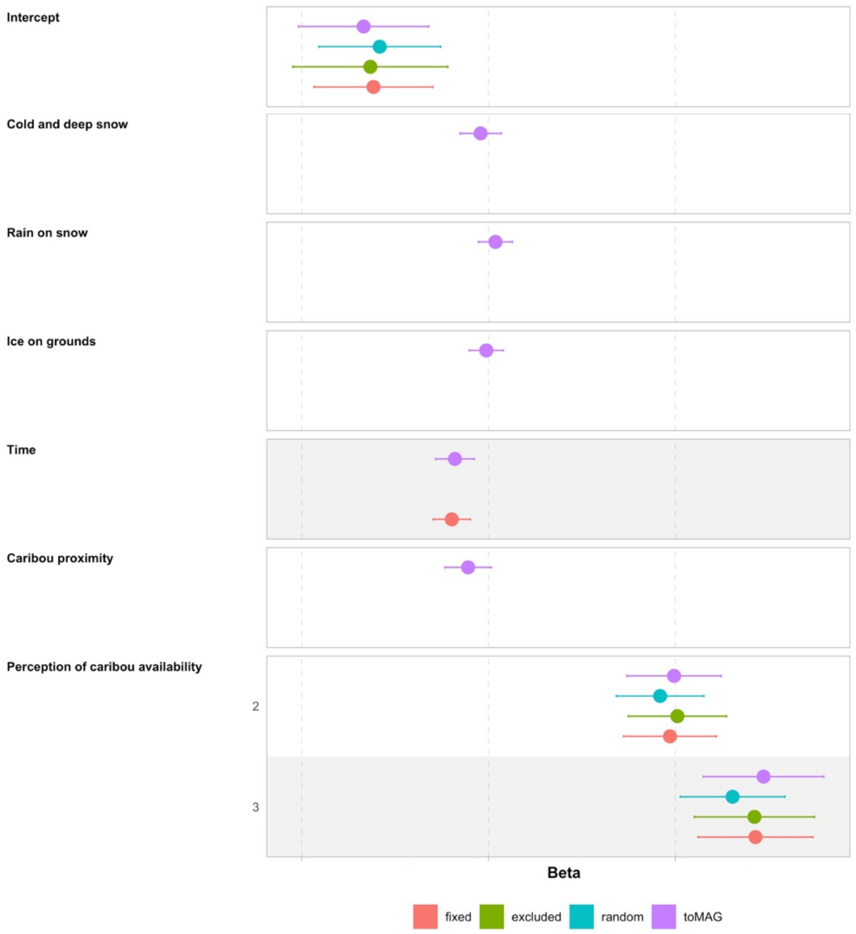

**Supplementary Fig. 5.** continued.

**d**. Effects on probability of meeting needs (step V, fall season)

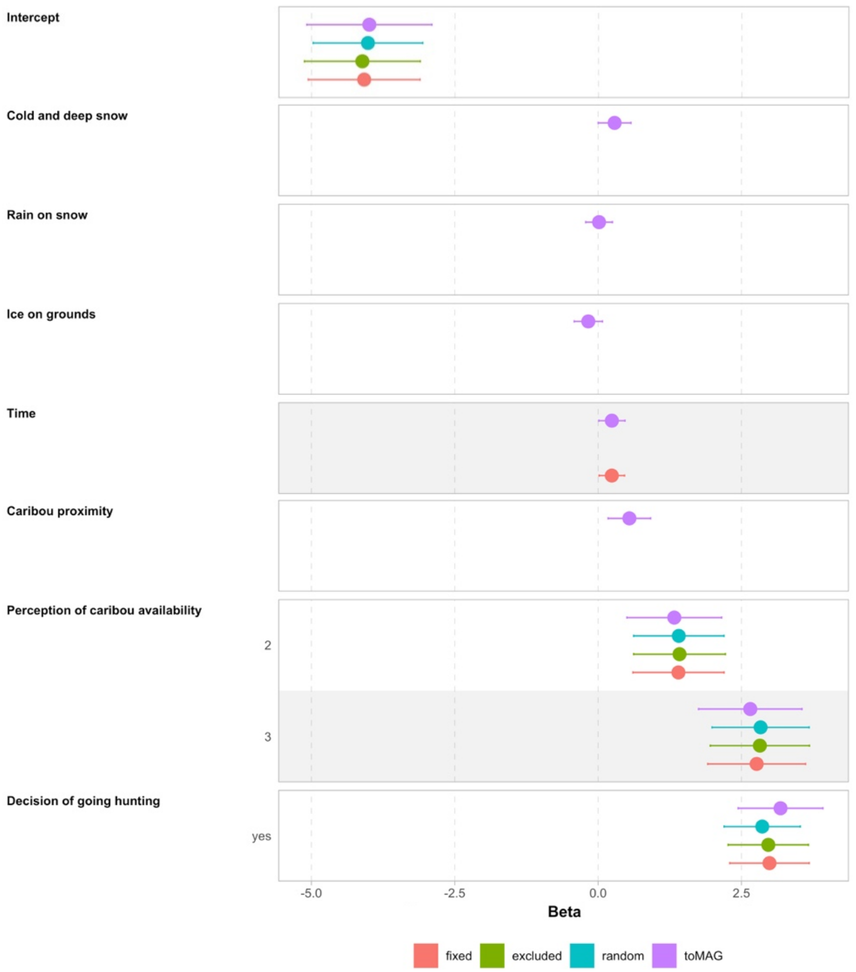

**Supplementary Fig. 5.** continued.

**a.** Effects on caribou proximity (step II, spring season)

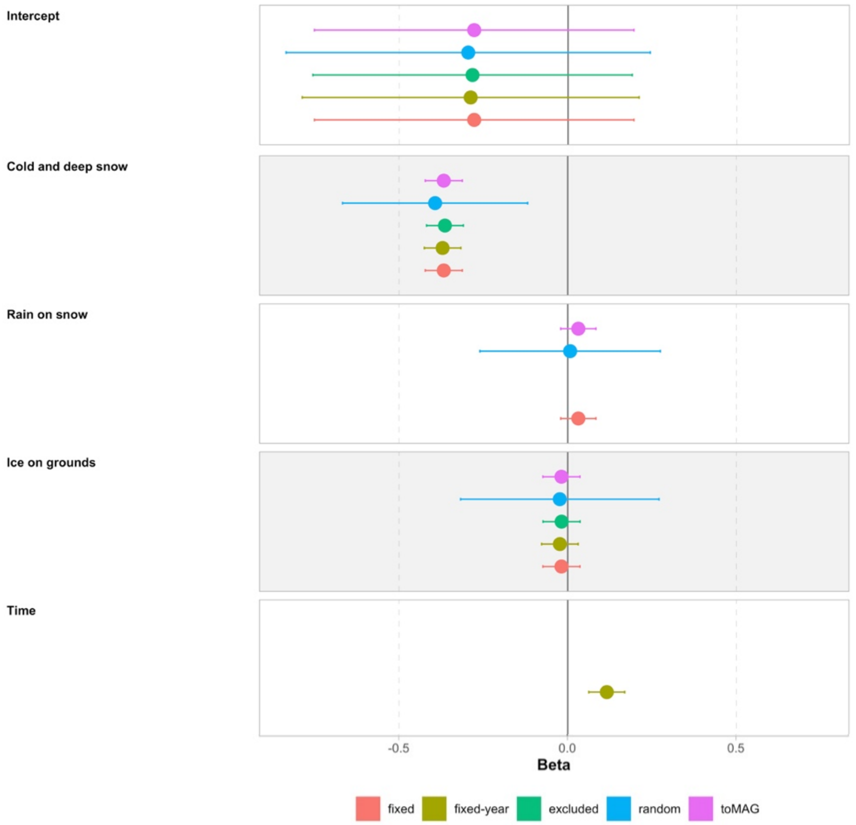

**Supplementary Fig. 6.** Mean parameter estimates (dots) and 95% confidence intervals (whiskers) contrasting five different methods to include the time effect "years" for the piecewise SEM of the spring season (n=616). The model presented in the main text is in red. This is a model including "temperature and early melt" as a fixed effect. This variable was strongly correlated with the variable "years" (time; r = 0.85), and therefore represented the combined effect of temporal changes and changes in snow and temperature conditions. In brown is the model including only the variable "years" (time) as a fixed effect and excluding the correlated variable "temperature and early melt". In green is the model fully excluding "years" and the correlated variable "temperature and early melt". In blue is the model including "years" as a random intercept. In purple is the model transforming the model including "years" as a fixed effect to a MAG.

**b**. Effects on perception of caribou availability (step III, spring season)

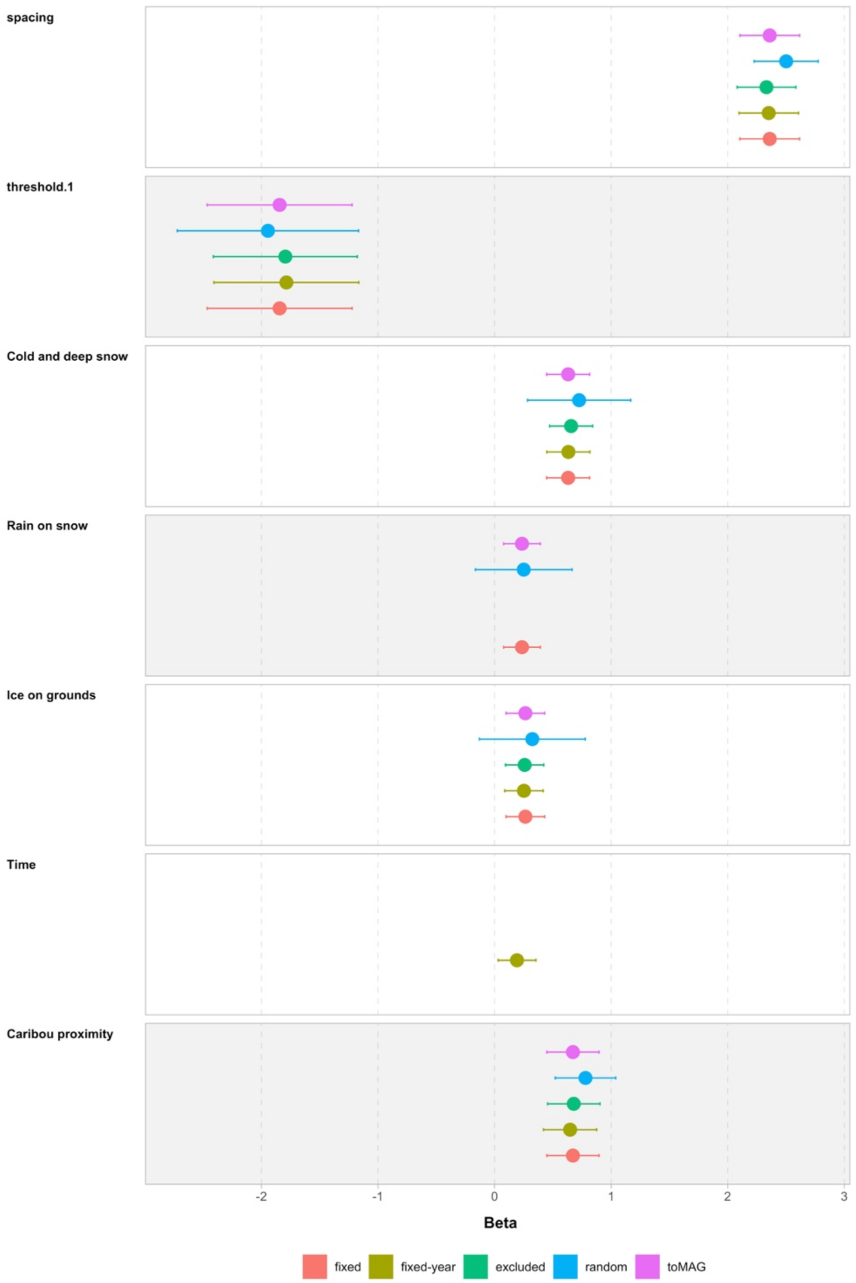

**Supplementary Fig. 6.** continued.

**c**. Effects on probability of going hunting (step IV, spring season)

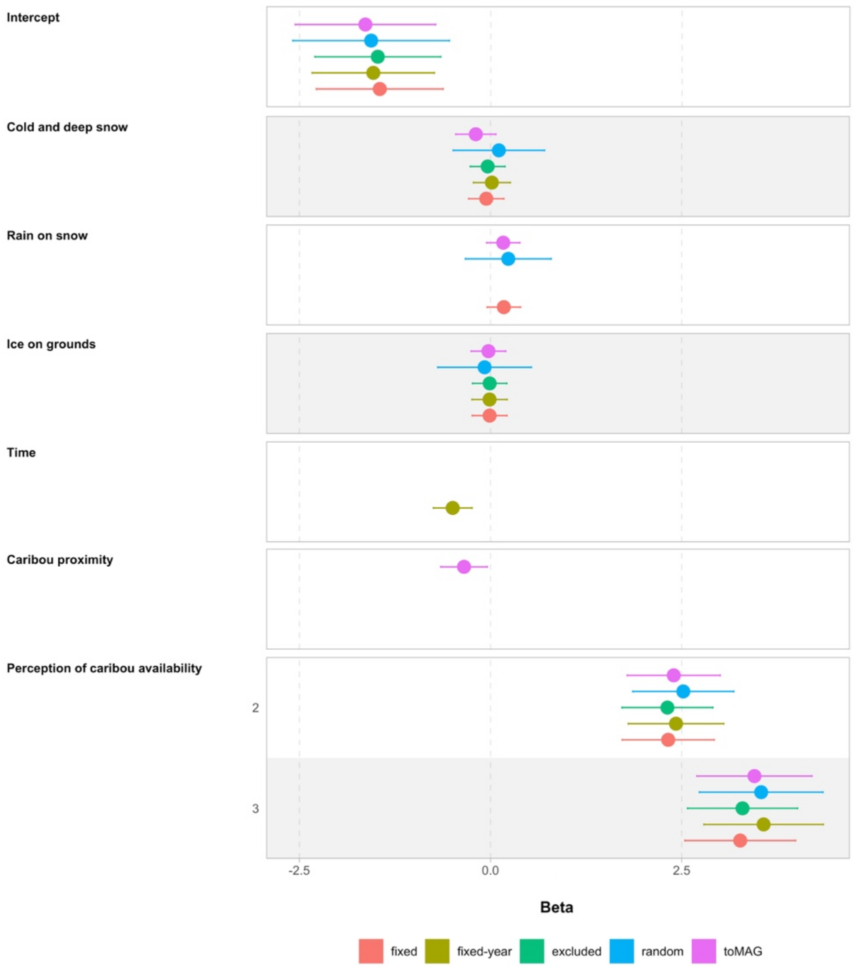

**Supplementary Fig. 6.** continued.

**d**. Effects on probability of meeting needs (step V, spring season)

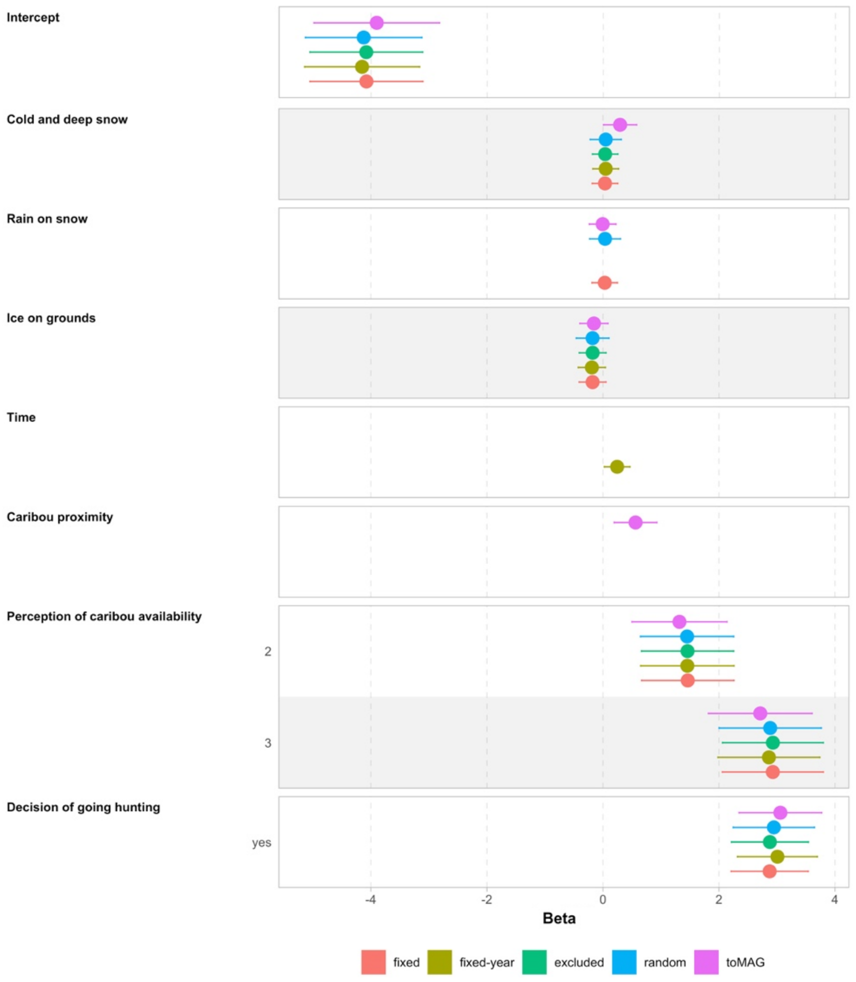

**Supplementary Fig. 6.** continued.

### Supplementary Tables 6 to 8

**Supplementary Table 6.** Complete list of potential competitive structural equation models evaluated for the influence of environmental conditions and time (years) on caribou distribution in relation to communities (proximity), hunter's perception of caribou availability, decisions to go hunting and of the capacity of hunters to meet their needs in caribou during fall.

| Models | ID* | NK | Cvalue | Pvalue | ΔAICc | $AICc_{weight}$ |
|---|---|---|---|---|---|---|
| STEP I) Causal structure among the exogenous variables, i.e. environmental conditions and time | | | | | | |
| **Null model** | **1a** | **0** | **4.76** | **0.97** | **0** | **0.92** |
| PCsnow ~ years, PCice1 ~ years, PCice2 ~ years | 1b | 6 | 3.00 | 0.81 | 52.24 | 0 |
| PCsnow ~ years | 1c | 2 | 4.43 | 0.93 | 5.66 | 0.05 |
| PCice1 ~ years, PCice2 ~ years | 1d | 4 | 3.17 | 0.92 | 16.41 | 0 |
| PCsnow ~ years, corr(PCsnow, PCice1) | 1e | 3 | 3.57 | 0.89 | 9.60 | 0.01 |
| PCsnow ~ years, corr(PCsnow, PCice2) | 1f | 3 | 2.54 | 0.96 | 8.57 | 0.01 |
| PCsnow ~ years, corr(PCsnow, PCice1), corr(PCsnow, PCice 2) | 1g | 4 | 1.82 | 0.94 | 15.05 | 0 |
| PCice1 ~ years, PCice2 ~ years, corr(PCsnow, PCice1) | 1h | 5 | 2.07 | 0.91 | 27.31 | 0 |
| PCice1 ~ years, PCice2 ~ years, corr(PCsnow, PCice2) | 1i | 5 | 0.85 | > 0.99 | 26.09 | 0 |
| PCice1 ~ years, PCice2 ~ years, corr(PCsnow, PCice1), corr(PCsnow, PCice2) | 1j | 6 | 0.14 | > 0.99 | 49.37 | 0 |
| STEP II) Effects of environmental conditions and time on CARIBOU PROXIMITY | | | | | | |
| **PCsnow + PCice1 + PCice2 + years** | **2a** | **6** | **4.76** | **0.97** | **0** | **1** |
| Null model | 2b | 2 | 230.82 | < 0.001 | 217.95 | 0 |
| PCsnow + PCice1 + PCice2 | 2c | 5 | 26.16 | 0.014 | 21.36 | 0 |
| years | 2d | 3 | 233.83 | < 0.001 | 222.98 | 0 |
| STEP III) Effects of environmental conditions, time and caribou proximity on PERCEPTION OF CARIBOU AVAILABILITY | | | | | | |
| **PCsnow + PCice1 + PCice2 + years + proximity** | **3a** | **15** | **4.76** | **0.97** | **0** | **0.98** |
| PCsnow + PCice1 + PCice2 + years | 3b | 14 | 37.88 | < 0.001 | 31.03 | 0 |
| Proximity | 3c | 11 | 137.20 | < 0.001 | 124.11 | 0 |
| PCsnow + PCice1 + PCice2 + proximity | 3d | 14 | 15.14 | 0.37 | 8.28 | 0.02 |
| years + proximity | 3e | 12 | 118.68 | <0.001 | 107.66 | 0 |
| STEP IV) Effects of environmental conditions, time and perception of caribou availability on HUNTING | | | | | | |
| PCsnow + PCice1 + PCice2 + years + availability | 4a | 24 | 9.73 | 0.78 | 4.40 | 0.10 |
| PCsnow + PCice1 + PCice2 + years | 4b | 22 | 70.95 | < 0.001 | 61.33 | 0 |
| availability | 4c | 20 | 33.23 | 0.059 | 19.35 | 0 |
| PCsnow + PCice1 + PCice2 + availability | 4d | 23 | 26.10 | 0.053 | 18.62 | 0 |
| **years + availability** | **4e** | **21** | **11.75** | **0.92** | **0** | **0.90** |
| STEP V) Effects of environmental conditions, time, perception of caribou availability and hunting on MEETING NEEDS | | | | | | |
| PCsnow + PCice1 + PCice2 + years + hunting | 5a | 29 | 47.03 | 0.003 | 21.23 | 0 |
| PCsnow + PCice1 + PCice2 + years | 5b | 28 | 152.55 | < 0.001 | 124.56 | 0 |
| hunting | 5c | 25 | 68.49 | < 0.001 | 34.01 | 0 |
| PCsnow + PCice1 + PCice2 + hunting | 5d | 28 | 61.43 | < 0.001 | 33.45 | 0 |
| years + hunting | 5e | 26 | 55.17 | 0.003 | 22.85 | 0 |
| PCsnow + PCice1 + PCice2 + years + availability + hunting | 5f | 31 | 22.84 | 0.41 | 1.42 | 0.30 |
| availability + hunting | 5g | 27 | 34.23 | 0.27 | 4.07 | 0.08 |
| PCsnow + PCice1 + PCice2 + availability + hunting | 5h | 30 | 30.04 | 0.18 | 6.43 | 0.02 |
| **years + availability + hunting** | **5i** | **28** | **27.98** | **0.47** | **0** | **0.60** |

*** Note:** At each step, the models include the effects selected in the previous steps. Models were selected at each step based on ΔAIC (difference in Akaike Information Criterion; see SI Materials and methods) and are identified in bold.

ID: model identification number, as used in the code. NK: the number of parameters in the model; Cvalue: Fisher's C statistic; Pvalue: The result of the Chi-squared distribution of the Cvalue with 2*k* degrees of freedom, where *k* is the number of independence claims; AICc: Akaike's Information Criterion corrected for small sample size; PCsnow: principal component on snow and temperature variables, contrasting years with colder temperatures, early snow arrival and more snow accumulation during fall versus years with warmer falls, late snow season and shallower snow conditions (see Methods and Supplementary Table 5); PCice1: first principal component on icing variables contrasting years with high frequencies and intensities of icing events (especially the amount of rain falling on snow) versus years with low frequencies and intensities of icing events; PCice2: second principal component on icing variables contrasting years with high versus low frequencies of freeze-thaw events and freezing rain falling directly on the ground.

**Supplementary Table 7.** Complete list of potential competitive structural equation models evaluated for the influence of environmental conditions and time[&] on caribou distribution in relation to communities (proximity), hunter's perception of caribou availability, decisions to go hunting and of the the capacity of hunters to meet their needs in caribou during spring.

| Models | ID* | NK | Cvalue | Pvalue | ΔAICc | $AICc_{weight}$ |
|---|---|---|---|---|---|---|
| STEP I) Causal structure among the exogenous variables, i.e. environmental conditions | | | | | | |
| **Null model** | **1a** | **0** | **2.65** | **0.85** | **0.00** | **0.31** |
| corr(PCice1, PCsnow1[years][&]) | 1b | 1 | 0.78 | 0.94 | 0.14 | 0.29 |
| corr(PCice1, PCsnow1[years][&]), corr(PCice1, PCsnow2) | 1c | 2 | 0.54 | 0.76 | 1.91 | 0.12 |
| corr(PCice1, PCsnow1[years][&]), corr(PCsnow1[years][&], PCsnow2) | 1d | 2 | 0.24 | 0.87 | 1.61 | 0.14 |
| corr(PCice1, PCsnow2), corr(PCsnow1[years][&], PCsnow2) | 1e | 2 | 1.33 | 0.51 | 2.70 | 0.08 |
| corr(PCice1, PCsnow1[years][&]), corr(PCice1, PCsnow2), corr(PCsnow1[years][&], PCsnow2) | 1f | 3 | 0 | 1 | 3.39 | 0.06 |
| STEP II) Effects of environmental conditions on CARIBOU PROXIMITY | | | | | | |
| Null model | 2a | 0 | 248.55 | 0 | 235.80 | 0.00 |
| **PCice1 + PCsnow1[years][&] + PCsnow2** | **2b** | **5** | **2.65** | **0.85** | **0.00** | **1.00** |
| PCsnow1[years][&] | 2c | 3 | 153.03 | 0 | 146.32 | 0.00 |
| STEP III) Effects of environmental conditions and caribou proximity on PERCEPTION OF CARIBOU AVAILABILITY | | | | | | |
| **PCice1 + PCsnow1[years][&] + PCsnow2 + proximity** | **3a** | **13** | **2.65** | **0.85** | **0.00** | **1.00** |
| PCice1 + PCsnow1[years][&] + PCsnow2 | 3b | 12 | 35.61 | 0.00 | 30.84 | 0.00 |
| Proximity | 3c | 10 | 71.34 | 0.00 | 62.42 | 0.00 |
| PCsnow1[years][&] | 3d | 11 | 39.29 | 0.30 | 32.45 | 0.00 |
| STEP IV) Effects of environmental conditions and perception of caribou availability on HUNTING | | | | | | |
| **PCice1 + PCsnow1[years][&] + PCsnow2 + availability** | **4a** | **21** | **3.54** | **0.90** | **0.00** | **0.99** |
| PCice1 + PCsnow1[years][&] + PCsnow2 | 4b | 19 | 50.48 | 0.00 | 42.59 | 0.00 |
| Availability | 4c | 18 | 31.32 | 0.01 | 21.39 | 0.00 |
| PCsnow1[years][&] + availability | 4d | 19 | 18.30 | 0.11 | 10.49 | 0.01 |
| STEP V) Effects of environmental conditions, perception of caribou availability and hunting on MEETING NEEDS | | | | | | |
| PCice1 + PCsnow1[years][&] + PCsnow2 + hunting | 5a | 28 | 45.90 | 0.00 | 34.71 | 0.00 |
| PCice1 + PCsnow1[years][&] + PCsnow2 | 5b | 27 | 120.71 | 0.00 | 107.32 | 0.00 |
| Hunting | 5c | 25 | 108.04 | 0.00 | 90.28 | 0.00 |
| PCsnow1[years][&] + hunting | 5d | 26 | 68.19 | 0.00 | 52.61 | 0.00 |
| **PCice1 + PCsnow1[years][&] + PCsnow2 + hunting + availability** | **5e** | **30** | **6.78** | **0.75** | **0.00** | **0.99** |
| PCsnow1[years][&] + hunting + availability | 5f | 28 | 20.23 | 0.10 | 9.72 | 0.01 |

*** Note:** At each step, the models include the effects selected in the previous steps. Models were selected at each step based on ΔAIC (difference in Akaike's Information Criterion; see SI Materials and methods) and are identified in bold.

[&] Since PCsnow1 was strongly correlated with years (time; r = 0.85), this variable represented the combined effect of temporal changes and changes in snow and temperature condition.

ID: model identification number, as used in the code. NK: the number of parameters in the model; Cvalue: Fisher's C statistic; Pvalue: The result of the Chi-squared distribution of the Cvalue with 2k degrees of freedom, where k is the number of independence claims; AICc: Akaike's Information Criterion corrected for small sample size; PCice1: first principal component on icing variables contrasting years with high versus low frequencies and intensities of icing events in general, and particularly the frequency of freeze-thaw events and the amount of freezing-rain falling directly on the ground during the winter (see Methods and Supplementary Table 5); PCsnow1: first principal component on snow variables describing years with shorter snow season in winter and spring, and higher temperatures during these seasons; PCsnow2: represents the second principal component on snow variables, contrasting years with deep snow and low variability in the snow cover versus years with shallow snow conditions and greater variability in the snow cover.

**Supplementary Table 8.** Values used to center and standardize the continuous variables used in the piecewise structural equation modelling (SEM).

| Variable | Season of analysis | Average | Standard deviation (SD) |
|---|---|---|---|
| PC1–Snow depth & Length of the cold snowing season | Fall | 0 | 1.48 |
| PC1–Frequency & intensity of all Icing events | Fall | 0 | 1.60 |
| PC2–Intensity of ground ice events (locked pastures) | Fall | 0 | 1.38 |
| Years | Fall | 2003.91 | 2.43 |
| Distribution in relation to communities[&] | Fall | 267.17 | 87.66 |
| PC1–Temperature & early melt * | Spring | 0 | 1.81 |
| PC2–Snow depth | Spring | 0 | 1.31 |
| PC1–Frequency & intensity of all Icing events | Spring | 0 | 1.68 |
| Distribution in relation to communities[&] | Spring | 260.89 | 77.82 |

* This PC is highly correlated with years (2000-2008; *r* [95% confidence interval] = 0.85 [0.44; 0.97]).

[&] Note that, for ease of interpretation with indices of caribou availability, we used in the analyses the median distribution to communities as the reverse of median distances.

### Supplementary Data and Code

The following datasets and codes are available via the Dryad permanent repository (https://doi.org/10.5061/dryad.msbcc2g2z)[29].

**Dataset S1 (separate file).** Database of climate variables likely affecting the Porcupine Caribou Herd (PCH) on its fall range.

**Dataset S2 (separate file).** Database of climate variables likely affecting the Porcupine Caribou Herd (PCH) on its spring range.

**Dataset S3 (separate file).** Database of median caribou distances (m) to communities for the spring and fall seasons.

**Code fall (separate file).** Rmarkdown (html) containing the R code and the results for the PCA and Path analysis performed for the fall season.

**Code spring (separate file).** Rmarkdown (html) containing the R code and the results for the PCA and Path analysis performed for the spring season.
